# Modeling Coupled Ecohydrological and Structural Responses to Forest Thinning

**DOI:** 10.64898/2026.11.13.766569

**Authors:** Wandi Feng, Benjamin Wong Blonder

## Abstract

Forest thinning is a silvicultural practice widely used to reduce competition among trees and enhance forest resilience. However, its long-term coupled effects on forest water balance and canopy structure remain uncertain due to competing hydrological processes, disturbances, and climatic variability. This study developed a multiyear stochastic ecohydrological model coupling soil saturation and water fluxes with canopy structure, represented by dynamic leaf area index (*LAI*), to examine how thinning alters soil water storage and evapotranspiration (*ET*) as the canopy regrows. Simulations were conducted for historically thinned forest sites, parameterized using FLUXNET climate data and species/stand-level traits obtained from the TRY database and published literature, with thinning intensity represented by initial *LAI* reductions ranging from 0% to 50%. Results show that long-term soil saturation responses and canopy recovery rates vary substantially across sites, while total *ET* remains relatively stable following thinning, as increased soil evaporation and greater water availability for the remaining canopy can compensate for reductions in stand-level transpiration. These findings highlight the importance of site-specific conditions in determining forest ecohydrological responses and the potential water conservation benefits of thinning.

## 2. Introduction

One central focus of modern forest management is achieving and maintaining an optimal stand density, neither so crowded that competition for space and resources suppresses individual tree growth, nor so sparse that site resources such as light, water, and nutrients are underused. Forest thinning is a widely used silvicultural practice for this purpose that selectively removes trees to improve forest resource-use efficiency while reducing vegetation stress (Hawthorne et al., 2013; Zeide, 2001). To adapt to different site conditions and management objectives, a range of thinning strategies has been used. In terms of structural selection, low thinning removes suppressed and intermediate trees with small crowns, reducing competition while retaining most of the upper-canopy structure. In contrast, crown thinning removes more dominant or codominant trees, thereby redistributing growing space among the remaining crown classes. Thinning treatments also vary in their spatial arrangement, with variable-density thinning creating a mosaic of areas with different residual tree densities. Thinning intensity, often expressed as the percentage of stems or basal area removed, also varies across treatments (Ashton & Kelty, 2017). Following thinning, vegetation can reoccupy released growing space through the lateral expansion of retained tree crowns and the establishment and growth of understory vegetation. Increased light availability can also slow lower-branch mortality and crown recession, helping maintain deeper live crowns and contributing to greater vertical canopy complexity (Bailey & Tappeiner, 1998; Chan et al., 2006; Dodson et al., 2012).

However, forest structure is highly complex and difficult to quantify, consisting of vertical layers, patches, gaps, and species heterogeneity that evolve dynamically through ecosystem processes such as growth, mortality, regeneration, and disturbance. Among all forest structural components, foliage is of particular functional importance because it is where the majority of light interception and gas and water exchange activities (photosynthesis, transpiration, and respiration) take place. One widely used way to characterize functional leaf space is through leaf area index (*LAI*), defined as the one-sided leaf area per unit ground surface area. *LAI* reduces the three-dimensional complexity of forest leaf structure to a tractable stand-level variable. In fact, *LAI* is a particularly useful proxy for forest growth because it well represents the occupancy of photosynthetic growing space and has been shown to strongly correlate with total volume increment at both the tree and stand levels (DeRose & Seymour, 2012). Thus, although forest growth occurs concurrently across multiple components, including roots, stems, and foliage, at differing rates and magnitudes, and with substantial structural heterogeneity across age and size classes, *LAI* remains a frequently used metric for characterizing forest growth, especially canopy development, at the stand level (Waring, 1983; DeRose & Seymour, 2012).

Following forest thinning, reduced light competition at the individual-tree level may promote growth and crown expansion through increased light availability, potentially enhancing light absorption and light-use efficiency. At the stand level, however, canopy removal generally decreases total light interception and may consequently reduce stand-level photosynthesis (Binkley et al., 2013; DeRose & Seymour, 2012). Besides light, water is another critical factor controlling forest photosynthetic capacity and health, serving as a central constraint on ecosystem function and productivity. One important objective of many thinning treatments is to augment forest water supply, and forest foliage is an important functional and structural factor controlling forest *ET* and throughfall. However, water-balance response to forest thinning involves both short-term and long-term tradeoffs. Immediately after thinning, the mechanism may appear straightforward: reduced leaf area (lower *LAI*) decreases stand-level transpiration and canopy interception. As a result, less water is lost through tree water use, and more precipitation reaches the soil as throughfall. At the same time, however, the canopy opening allows more sunlight and wind to enter the stand, which can increase soil and understory *ET*. Moreover, transpiration itself also responds to both light and water availability. Greater light availability can increase photosynthetic demand, promoting stomatal opening so that carbon dioxide can enter the leaf; however, open stomata also allow water vapor to exit through transpiration (Sohn et al., 2016; Bian et al., 2026). Soil water availability further regulates transpiration by constraining water supply to the leaves and interacting with plant water-use strategies, particularly stomatal regulation in response to water stress (Bassiouni et al., 2023). Therefore, although total stand-level transpiration can decrease after thinning, at the individual tree level, transpiration may actually increase due to reduced competition for water. In the long term, as retained trees expand their crowns and ground vegetation develops, transpiration and canopy interception can increase again, which may reduce, or even reverse, the initial water benefit of thinning (Sohn et al., 2016; Bian et al., 2026).

Although many field studies have examined thinned forest sites, few have been able to track the long-term effects of thinning on forest ecohydrological dynamics, and isolating the effects of thinning itself remains difficult given interannual climate variability and environmental disturbances. To address this, an ecohydrological model is developed to couple forest foliage growth, represented by changes in *LAI*, with forest water fluxes, including *ET*, using soil saturation as the linking state variable. With this framework, the study aims to examine: (1) How does forest thinning affect long-term forest water storage (i.e, soil saturation)? (2) How does thinning alter total *ET* and its partitioning between soil evaporation and tree transpiration, and how do these effects change during canopy recovery? (3) How does the effect of thinning vary across plant functional types and site climate and soil conditions?

## 3. Methods

To study multi-year forest water dynamics and canopy recovery, this study develops a modeling framework coupling growing-season ecohydrological processes with annual leaf area growth. At the growing-season scale, the framework consists of two components. First, building on the ecohydrological model of Bassiouni et al. (2023), which incorporates the soil–plant–atmosphere continuum (SPAC) to describe water transport from the soil through plants to the atmosphere, the model calculates steady-state soil saturation probability distributions and associated *ET* fluxes. Second, a modified formulation based on Vico & Porporato (2013) is used to estimate growing-season leaf area growth (increase in *LAI*) constrained by both soil saturation and light interception. At the annual scale, *LAI* is updated based on the net balance between growing-season leaf area increment and annual leaf turnover, with the updated *LAI* feeding back into the following year’s ecohydrological calculations. The model is parameterized using FLUXNET climatic data and site/species-specific characteristics obtained from published field studies of selected historically thinned forests to simulate thinning responses under different environmental conditions.

### 3.1 Growing-Season Ecohydrology and Leaf Growth

#### 3.1.1 Stochastic Water Balance Equation

Following Laio et al. (2001), the instantaneous rate of change of soil saturation *s*(*t*) ∈ [0, 1] is defined by the stochastic water balance at a point:

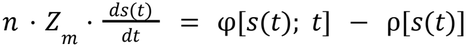

where *n* (*m*^3^ *m*^−3^) is soil porosity and *Z*_*m*_ (*m*) is the rooting zone depth. The left-hand side represents the rate of change of water stored per unit area within the active rooting zone.

The input function φ[*s*(*t*); *t*] describes the rate of water infiltrating into the soil column:

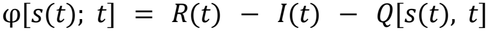

where *R*(*t*) is the rainfall rate, modelled as a Poisson process with mean event frequency λ (*day*^−1^) and exponentially distributed event depths with mean α(*m*); *I*(*t*) is the rainfall depth lost to canopy interception prior to reaching the soil surface; *Q*[*s*(*t*), *t*] is surface runoff in which the excess depth is instantaneously removed whenever an incoming rainfall event drives *s*(*t*) above unity.

The loss function ρ[*s*(*t*)] describes the deterministic state-dependent rate of water leaving the rooting zone:

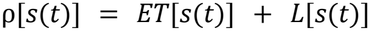

where *ET*[*s*(*t*)] is the *ET* rate and *L*[*s*(*t*)] is the leakage rate representing gravitational drainage below the rooting zone, which is only non-zero when *s*(*t*) exceeds field capacity (*s*_*fc*_).

Here, all components of the water balance function are expressed in units of *m day*^−1^.

#### 3.1.2 Water Loss Function

Extending Laio et al. (2001), Bassiouni et al. (2023) derive the loss function ρ[s(t)] by first partitioning the total potential evapotranspiration *E*_0_ (*m day*) ^−1^ between soil evaporation and plant transpiration using Beer-Lambert law. The fraction of incoming radiation penetrating through the canopy to the soil surface is given by:

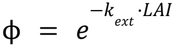

where *k*_*ext*_ is the canopy light extinction coefficient. This partitioning yields two normalised water loss rates (*day*^−1^), each expressed as a fraction of the total soil water storage capacity *n* · *Z*_*m*_ :

Maximum soil evaporation: 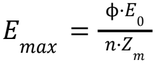

Well watered plant transpiration: 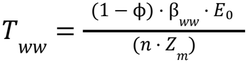

Here β_*ww*_ is the well-watered transpiration ratio, the fraction of E_0_ that the plant can sustain under unlimited soil water supply, derived analytically from the non-dimensional Π groups characterising plant hydraulic traits and atmospheric demand (A.1.2.2; Bassiouni et al., 2023)

Normalized by the total soil water storage capacity *n*·*Z*_*m*_, the loss function ρ[*s*(*t*)], now with units of *day*^−1^, is expressed as the following piecewise continuous function:

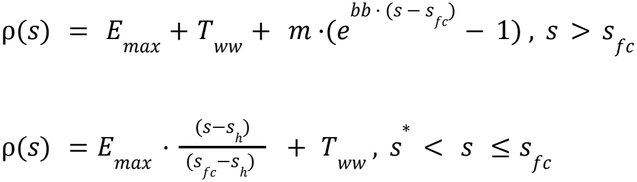

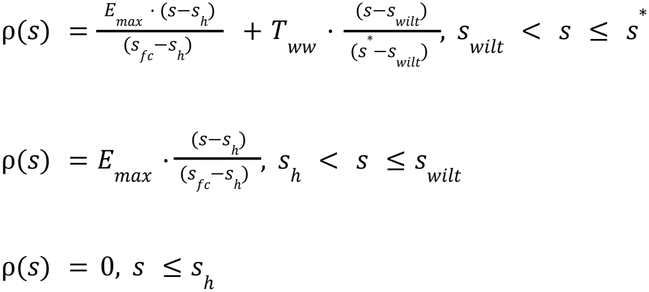

The soil evaporation component increases linearly from zero to *E*_*max*_ between the hygroscopic point *s*_*h*_, below which water is bound to soil particles and unavailable for evaporation, and field capacity *s*_*fc*_ (Fig.1.1). The plant transpiration component increases linearly from zero to *T*_*ww*_ between the wilting point *s*_*wilt*_ and the point of incipient stomatal closure *s*\* (Fig.1.1). Beyond *s*_*fc*_, ρ(*s*) additionally accounts for leakage/drainage which increases exponentially with *s. s*_*h*_ and *s*_*fc*_ are soil-texture constants determined by the soil water retention curve (A.2.1). *s*_*wilt*_ and *s*\* are derived from plant hydraulic traits by inverting the SPAC model to find soil water potential at which transpiration reaches prescribed fractions of the well watered ratio β_*ww*_ (5% for *s*_*wilt*_ and 95% for *s*\*), then converting that potential to soil saturation using the Brooks-Corey relationship (A.1.2.3; Bassiouni et al., 2023).

**Figure 1.1.**
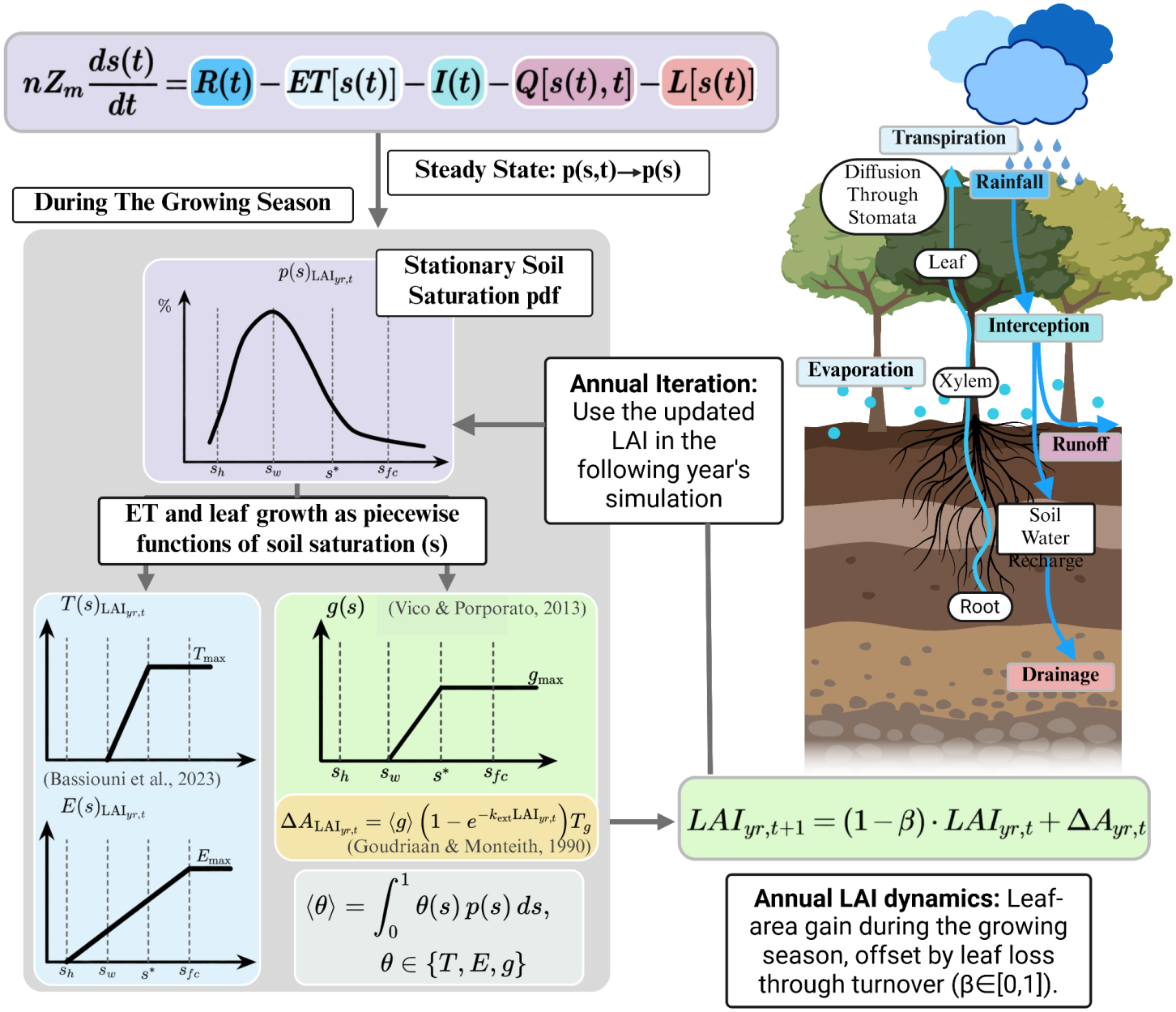
Schematic illustration of the model structure and forest water balance.

As β_*ww*_ inherently encodes plant physiological traits — including the water potential at 50% stomatal conductance loss Ψ_*g50*_ (*MPa*) (stomatal sensitivity) and the water potential at 50% xylem conductance loss Ψ_*x50*_ (*MPa*) (xylem vulnerability), the separation between *s*_*wilt*_ and *s*\*, which defines the soil saturation range over which the plant progressively closes its stomata in response to declining water supply, reflects the plant’s water use strategy in balancing carbon assimilation against the risk of hydraulic damage (Bassiouni et al., 2023).

#### 3.1.3 Analytical Soil Moisture Distribution and Seasonal Water Balance Components

The probability density function of soil moisture *p*(*s*) is thus solved analytically assuming growing season steady state (A.1.2.6; Bassiouni et al., 2023). Mean growing-season soil evaporation < *E* >, and tree transpiration, < *T* >, both expressed in *m day* ^−1^, are then calculated by integrating their respective soil saturation dependent loss function over this distribution:

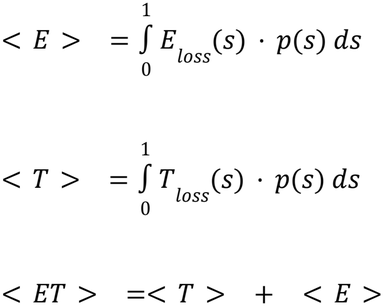

(A.1.2.7; Bassiouni et al., 2023)

#### 3.1.4 Growing Season Vegetation Growth Dynamics

Following the framework of Vico & Porporato (2013), the instantaneous growth rate *g*[*s*(*t*)] is defined as a piecewise, continuous function of relative soil saturation *s*(*t*):

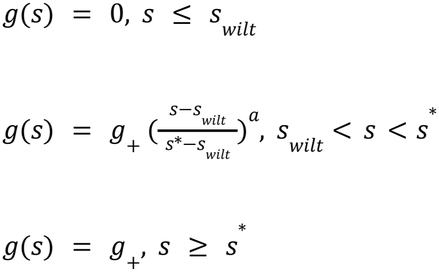

where *g*_+_ is the maximum, well-watered growth rate and *a* is a sensitivity parameter governing how strongly vegetation growth declines with decreasing soil saturation. The two thresholds are deliberately chosen to match the same thresholds used in the transpiration function of the soil water balance (Fig.1.1).

The mean growth rate < *g* > is then obtained numerically by integrating *g*(*s*) against the steady-state soil saturation pdf *p*(*s*):

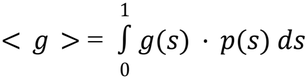

< *g* > inherits its units directly from *g*_+_. Whereas the original Vico & Porporato (2013) formulation was developed to describe crop biomass accumulation, here, to match the study’s focus on *LAI* and the daily timestep of the soil water balance model, *g*_+_ (and hence < *g* >) is instead interpreted as a daily increase in *LAI* per unit ground area (*m*^2^ *m*^−2^ *day*^−1^).

### 3.2 Interannual *LAI* Dynamics

The ecohydrological models discussed previously, including Laio et al. (2001), Vico and Porporato (2013), and Bassiouni et al. (2023), primarily describe within-growing-season soil moisture and plant water-use dynamics. To address the long term coupling between leaf area dynamics and forest ecohydrology after thinning, the present model thus links successive years by treating *LAI* as an annually updated state variable. In each year, leaf-area growth is derived from the steady-state ecohydrological conditions. The resulting *LAI* is then used to parameterize the within-growing-season soil-moisture dynamics for the following year, producing updated estimates of growth and water-flux rates (Fig.1.1).

From the previous section, we obtained< *g* > (*m*^2^ *m* ^−2^ *day*^−1^) as the mean daily leaf-area expansion rate per unit ground area. This rate is limited only by water availability. However, as discussed in the Introduction, photosynthetic capacity, and therefore foliage expansion, is also strongly constrained by canopy light interception. Because the model assumes spatially uniform *LAI* and represents the canopy as vertically aggregated and horizontally homogeneous, light limitation can be incorporated using a simple Beer–Lambert interception fraction. Assuming that *g*_+_ represents the well-watered growth rate under complete light interception, the growing-season leaf-area increment is scaled by the fraction of radiation intercepted by the canopy, 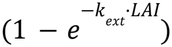, consistent with the formulation used above for *ET* partitioning 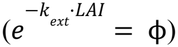 (Goudriaan & Monteith, 1990).

Marginal light interception per unit *LAI* declines exponentially as the canopy fills: each leaf layer absorbs a fixed fraction of the light reaching it, upper leaves intercept full incoming radiation while lower layers receive progressively attenuated light, producing a saturating relationship between *LAI* and light absorption in which each additional unit of leaf area yields diminishing returns (Binkley et al., 2013). This saturation also has implications for productivity: as *LAI* increases, lower leaves may become increasingly shaded and contribute little to whole-canopy carbon gain, or even become carbon-negative when photosynthesis no longer offsets respiratory costs as light availability falls below the leaf light compensation point. This effect is especially pronounced when the light extinction coefficient is high, as occurs in denser, more horizontally oriented, or more strongly clumped leaf layers. (Monsi and Saeki, 2005).

Scaled by the length of the growing season (*T*_*g*_ (*day*)), the total growing-season leaf-area increment can therefore be written as:

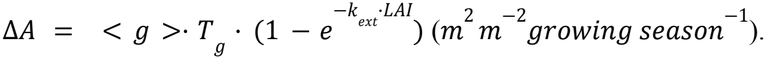

For deciduous trees, photosynthesis is strongly seasonal, with minimal photosynthetic activity during winter because the leaves are shed in autumn. In evergreen trees, leaves or needles are retained year-round, so low levels of photosynthesis may continue during mild winter periods. However, new shoots and needles generally develop after spring bud break, while winter is typically a dormant period with little active shoot growth. Therefore, annual foliage growth can be approximated as occurring primarily during the growing season with the timing and length of the growing season varying among species and geographic locations (Flint, 1974).

Not all leaves are retained into the following year as in a carbon balance scale, as not all assimilated carbon by photosynthesis turned into new biomass as some is lost through respiration and the net change in biomass also includes loss of existing tissues through litterfall. To describe annual changes in leaf area, we account for both new leaf production and leaf loss through shedding, yielding the following annual balance:

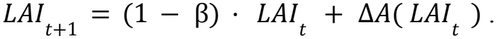

Where β is the fraction of foliage lost annually. At an annual time scale, the model aggregates the seasonal rise and fall of leaf area into annual fractions of foliage production and litterfall. This formulation can be applied to both deciduous and evergreen trees. For example, in deciduous forests, the net annual change in *LAI* may be zero even though substantial leaf production and loss occur within the year (Mäkelä & Valentine, 2020).

It is important to note that, unlike the growth term, β is treated here as a constant. This rate may be derived from field measurements or represented using leaf-age cohorts (Mäkelä & Valentine, 2020). However, litterfall and additional water-stress-induced senescence may vary with soil-moisture conditions, thereby linking foliage loss back to the ecohydrological model. A limitation is that the steady-state assumption is likely valid only within the growing season because of seasonal climatic variability. Litterfall may occur year-round, as in some evergreen forests, or primarily after the growing season. Therefore, estimating a soil-saturation-dependent litterfall rate would likely require a seasonal ecohydrological model.

**Figure 1.2.**
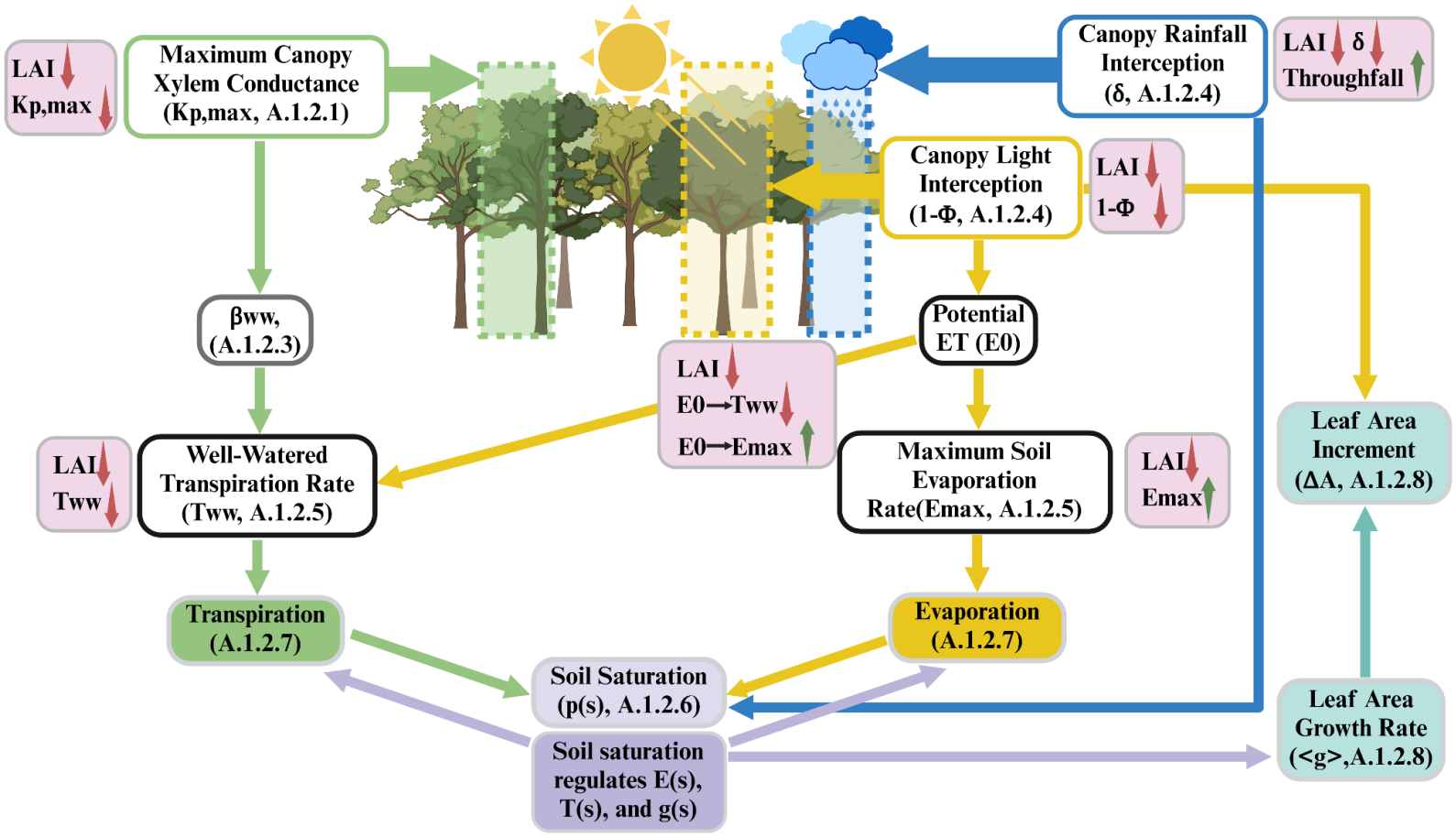
Schematic illustration of feedbacks between forest canopy leaf area and ecohydrological processes. *LAI* enters the model through three canopy-scale processes: (1) canopy conductance capacity, which regulates transpiration (green); (2) light interception, which partitions *E*_0_ between *T*_*ww*_ and *E*_*max*_, while also controlling leaf area growth (yellow); and (3) rainfall interception, which determines throughfall reaching the soil (blue). The boxes represent affected model variables, with arrows indicating the connecting pathways, while the pink annotations illustrate the direction of change in variables following *LAI* reduction. Together, these processes act on soil saturation (purple), which in turn regulates *ET* and leaf area growth, establishing feedback between canopy structure and ecohydrological dynamics.

### 3.3 Model Parameterization and Simulation

For model calibration and testing, four forest sites were selected for simulation. These sites were either historically thinned or had undergone regular thinning management. The sites were selected based on (1) the availability and quality of FLUXNET meteorological data, (2) the presence of a clearly dominant tree species and relatively simple stand structures, and (3) a well-defined primary growing season. Three of the four selected study sites are pine-dominated coniferous forests, which are commonly subjected to thinning, particularly in North America. Many tropical forests (high species diversity, complex vertical canopy structures, and potentially year-round growth), as well as Mediterranean forests exhibiting pronounced bimodal growth patterns, were considered less suitable for the simplifying assumptions of this modeling framework. Evergreen forests were also preferred because the model uses a single representative *LAI* value for each growing season. Although evergreen trees produce new foliage during the growing season, the stand-level *LAI* generally remains relatively stable. In contrast, deciduous forests experience substantial within-season changes in *LAI*, which can lead to corresponding changes in soil water availability and *ET* (Denham et al., 2023). Therefore, the model’s aggregated growing-season representation of coupled LAI and ecohydrological dynamics may be less suitable for deciduous forests.

It is important to note that the simulations are not intended to reproduce the exact observed response of each site or to make precise year-by-year predictions. The model is deliberately simplified and therefore cannot capture all processes that influence post-thinning hydrological and vegetational dynamics. Instead, its primary goal is to isolate the coupled interactions among selected factors, particularly leaf area, soil saturation, and water fluxes (*ET, T, E*).

Using site-specific data provides realistic and internally consistent combinations of vegetational, climate, and environmental parameters with which to examine the model. The results should therefore be interpreted as comparative experiments rather than exact forecasts. Simulations across different thinning intensities, initial stand structures, and climatic conditions can reveal how these factors shape the magnitude and persistence of thinning responses. Comparisons with existing field observations from the same sites can then be used to assess whether the model reproduces the main qualitative patterns and mechanisms observed in real forests, rather than requiring exact numerical agreement.

#### 3.3.1 Sites background and thinning history

Sites are identified by their FLUXNET tower station labels. The four selected sites are US-Fmf (Flagstaff Managed Forest), CA-TP4 (Turkey Point 1939 Plantation), FR-Pue (Puéchabon State Forest), and FI-Hyy (Hyytiälä SMEAR II boreal forest). Two of these sites, US-Fmf and CA-TP4, are temperate evergreen pine forests; FR-Pue is a Mediterranean evergreen oak forest; and FI-Hyy is a boreal evergreen pine forest.

All four sites have a history of thinning, but the timing and intensity of thinning differ among sites. The US-Fmf site underwent substantial thinning in September 2006, during which tree density was reduced by 70% and forest *LAI* was reduced by 40% (Dore et al., 2012). The CA-TP4 site was thinned in February–March 2012, with approximately one-third of the trees commercially harvested and *LAI* reduced by about 38% (Skubel et al., 2015). The FR-Pue site was thinned in March 1986 using selective thinning from below, in which smaller stems were preferentially removed. Experimental plots were assigned basal-area removal treatments of 0% (control), 25%, 45%, 60%, and 80%. Based on *LAI* estimated from diameter at breast height (DBH), these treatments corresponded approximately to immediate *LAI* reductions of 0%, 20%, 40%, 60%, and 77%, respectively (Cabon et al., 2018). FR-Pue also underwent an experimental thinning and rainfall-exclusion treatment in the winter of 2002–2003, during which approximately 30% of the stand basal area was removed from the thinned plots (Gavinet et al., 2019). The FI-Hyy site underwent understory removal in 2019 followed by a main thinning in February–March 2020. The treatment reduced basal area by approximately 40%, while *LAI* was reduced by about 40–45% (Aslan et al., 2024; Peräkylä et al., 2025).

#### 3.3.2 Climate inputs

Annual leaf area growth is assumed to occur entirely within the growing season. Here, the growing season is specifically defined as the period of active leaf area development. For deciduous species, this corresponds to the period between leaf emergence and fall coloration. For evergreen species, because leaves persist across multiple years, bud break and the initiation of new growth can be treated as phenological markers of the start of the growing season, while the end is indicated by the cessation of shoot growth after leaf area reaches its seasonal maximum (Lieth, 1974). Here, as all species included in this simulation are evergreen, site-specific growing-season lengths are obtained from literature reports (A.2) describing the period between bud break and the attainment of maximum *LAI*.

All climatic inputs are obtained from FLUXNET data and aggregated to daily time steps. Rainfall frequency (λ) and mean rainfall event depth (α) are derived directly from growing-season precipitation records. Average potential evapotranspiration (*E*_0_) is calculated using the Priestley–Taylor method implemented in the R package *Evapotranspiration* (Guo et al., 2016). Because relative humidity is not directly available in the FLUXNET dataset, it is calculated from air temperature and vapor pressure deficit (*VPD*) before being used in the *E*_0_ calculation with the R package *plantecophys* (Duursma, 2015).

Here, precipitation and *E*_0_ are calculated using growing-season data averaged across all years with complete FLUXNET records, regardless of when thinning treatments occurred. These growing-season climatic parameters are then held constant across simulated years, assuming stationary interannual climatic conditions to isolate the effects of thinning and subsequent canopy recovery. Additionally, for earlier thinning treatments, such as those conducted at FR-Pue in 1986, FLUXNET data from the treatment period are not available. As emphasized previously, the major purpose of using site-specific data in these simulations is not to reproduce or directly compare model predictions with observed responses to historical thinning events, but rather to establish realistic and internally consistent parameterizations.

#### 3.3.3 Soil property inputs

Soil is categorized into texture classes based on the relative proportions of sand, silt, and clay, as determined from literature describing the study sites (A.2). Following the same parameterization framework as Bassiouni et al. (2023), soil property inputs (*n, K*_*s*_, *b*, and Ψ_*s*0_) are assigned texture-specific values from Rawls et al. (1982). The hygroscopic point *s*_*h*_ is calculated from the soil-water retention curve as the soil saturation corresponding to a water potential of −10 *MPa*, and field capacity *s*_*fc*_ is taken as the saturation corresponding to −0.03 *MPa* (A.2.1).

Here, *Z*_*m*_, the effective soil depth, is defined as the layer of soil participating in the modeled water balance by actively responding to precipitation recharge and *ET* losses. It can therefore be interpreted as an effective rooting-zone depth. Because this study focuses primarily on foliage structure and field measurements of rooting depth are not consistently available across sites, a default value of *Z*_*m*_ is used. Laio et al. (2001) examined model sensitivity to *Z* over a range of 0.3–0.9 m and showed that shallower soil depths produce faster wetting and drying and therefore larger fluctuations in soil moisture with a broader *p*(*s*) distribution and greater sensitivity to drought, while mean soil moisture is comparatively less affected by changes in *Z*_*m*_. For consistency and simplicity, *Z*_*m*_ is therefore set to 0.45 m for all sites. This value is broadly consistent with global estimates of effective rooting-zone depth for where the four sites are located and is representative of the near-surface rooting zone within which soil-moisture variations are strongly coupled to *ET*, although the exact active depth varies with vegetation type and site-specific soil conditions (Yang et al., 2016; A et al., 2019).

#### 3.3.4 Vegetation inputs

Pre-thinning initial leaf area index (*LAI*_0_) and canopy height (*h*_*c*_) are derived either from FLUXNET BIF data or from literature-reported field measurements (A.2). Due to the lack of consistent field measurements across sites, root area index (*RAI*) values are assigned based on biome-level global averages reported by Jackson et al. (1997).

Similarly, plant hydraulic traits (Ψ_*g*50_, Ψ_*x*50_, and *k*_*xl,max*_) are also not consistently reported in the site-specific field studies. Therefore, default values were assigned using mean trait values from a dataset compiled using the TRY plant trait database (Kattge et al., 2020), grouped by climate and plant functional type (US-Fmf and CA-TP4: temperate gymnosperms, FR-Pue: Mediterranean evergreen angiosperms, FI-Hyy: boreal gymnosperms) (A.2).

#### 3.3.5 Maximum growth rate and growth sensitivity exponent

*g*_+_ as the maximum daily increase in *LAI* per unit ground area is not generally a field-measurable value. Therefore, the input value of *g*_+_ is numerically back-calculated such that β · *LAI*_*unthinned, t*_ = Δ*A*(*LAI*_*unthinned, t*_). This assumes that, for the unthinned stand and the specified input value of *g*_+_, the annual leaf area increment perfectly offsets litterfall, such that *LAI* remains stationary across years.

For the Flagstaff forest (US-Fmf), *LAI* measurements on the control site from 2006 (the year of thinning) to 2010 show little to no change in *LAI* (Dore et al., 2012). CA-TP4 is a white pine plantation established in 1939, which was considered mature when it was thinned in 2012, and *LAI* measurements from 2005 and 2011 show no strong change (peak of 8.0 vs. 8.6) (Chen et al., 2006; Wang & Chen, 2012). At FR-Pue, the experimental stands were 44–59 years old since the last clear-cut at the time of thinning, and a subsequent modeling study explicitly assumed that *LAI* in undisturbed stands remained constant over time (Cabon et al., 2018). Similarly, the FI-Hyy site is considered a 60-year-old boreal forest with a fully developed canopy structure (Aslan et al., 2024). Based on the above, it is generally reasonable, at least for these study sites, to assume that *LAI* remains stationary in unthinned control scenarios.

Setting the unthinned stand to a stationary *LAI* makes the pre-thinning *LAI*_0_ an equilibrium of the model, toward which the simulated post-thinning *LAI* trajectories recover. Because *LAI* is the only dynamic variable in the simulation, the other outputs, such as soil saturation and water fluxes, also return to the same pre-thinning equilibrium as *LAI* recovers. This co-return of canopy structure and hydrological conditions is consistent with the theory by Eagleson (1982) that natural soil–vegetation systems tend toward a stable equilibrium with their climatic and edaphic environments.

The growth sensitivity exponent (*a*) is set to 1 in this simulation, resulting in a linear relationship between leaf area growth rate and soil saturation within the water-limited range (*s*_*wilt*_ < *s* < *s* *). A higher exponent *a* = 6 was used in Laio et al. (2001), whereas a similar piecewise-linear growth response to relative soil water content (*a* = 1) was used to model tree-ring growth by Evans et al. (2006).

Here, *a* = 1 was chosen so that the growth function has the same piecewise-linear form as the transpiration function.

#### 3.3.6 Simulation scenario settings

For each site, simulations are run under six different thinning intensities, represented as percentage reductions in initial *LAI*: 0% (control), 10%, 20%, 30%, 40%, and 50%. Although thinning can be conducted at higher intensities in practice, this model does not explicitly consider regeneration as it would make it difficult to define a stand-level uniform growth rate. Depending on tree age, compensatory growth of residual trees has limits at the stand-level (Sun et al., 2026). For example, if tree-density removal is assumed to be proportional to *LAI* reduction, then a 50% reduction in tree density (assuming a corresponding 50% reduction in *LAI*) would require the remaining trees to support approximately twice as much leaf area as before for the stand to fully recover its initial *LAI*. Because the model treats the canopy as a spatially uniform whole and does not include a mechanism that constrains the maximum leaf area that an individual tree can support, simulations at high thinning intensities should therefore be interpreted with caution.

For each site, soil saturation, water fluxes, and leaf area trajectories are simulated over a 50-year period under each thinning intensity.

**Figure 1.3.**
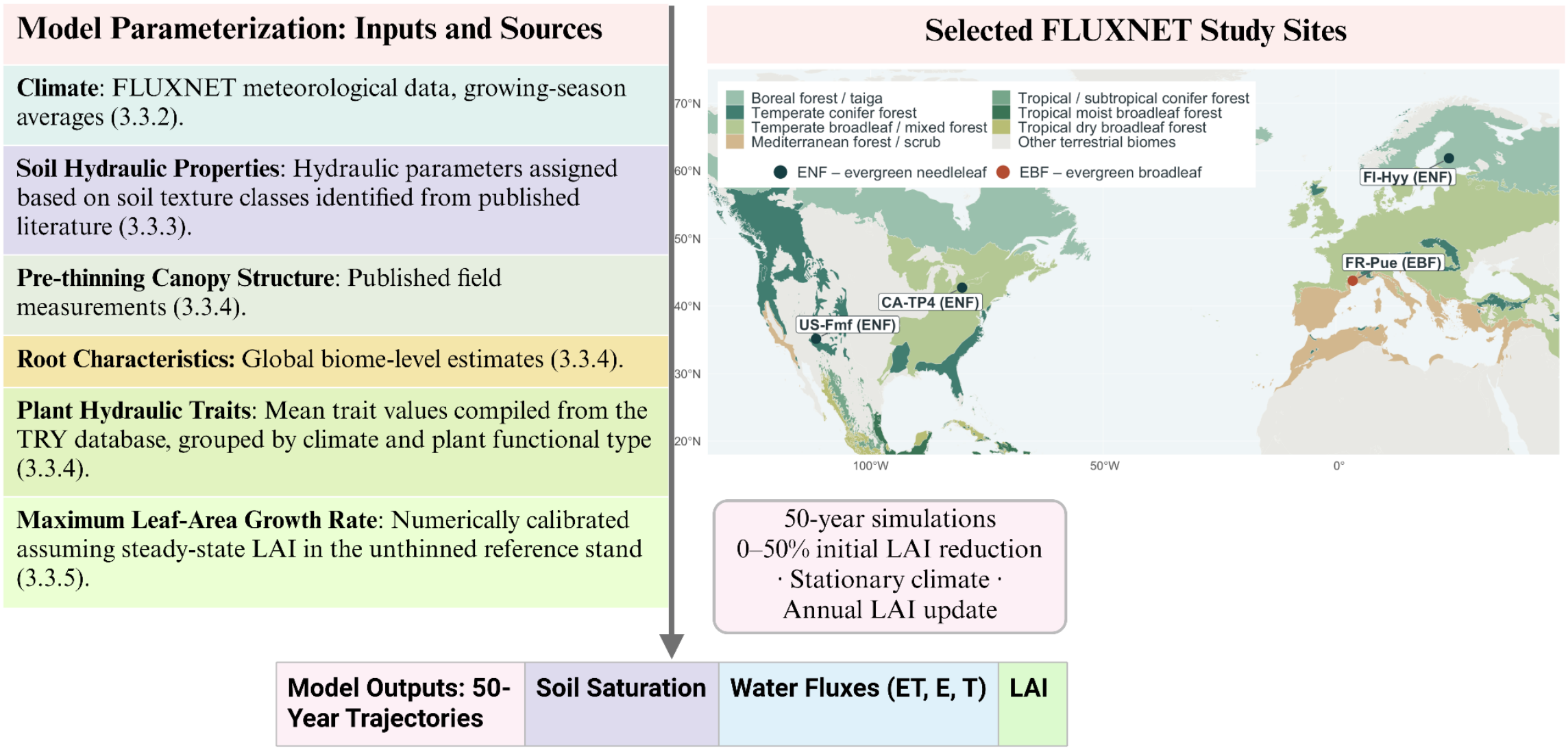
Model parameterization and simulation workflow.

### 3.4 Sensitivity Analysis

To assess the sensitivity of model outputs to key inputs representing soil and root properties, plant hydraulic traits, and climatic forcing, a local elasticity analysis was performed for three categories of parameters: (1) soil and root properties (*Z*_*m*_ & *RAI*); (2) plant hydraulics (Ψ_*g50*_, Ψ_*x50*_, *k*_*xl,max*_); and (3) climatic inputs (α, λ, *E*_0_). For selected thinning scenarios (0%, 30%, and 50%), we calculated the elasticity of first-year model outputs relative to changes in each input parameter using a central finite-difference approximation. The elasticity was calculated as follows:

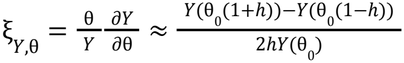

where *Y* ∈ {< *s* >, < *T* >, < *E* >, < *ET* >} represents the model outputs, θ ∈ {*Z*_*m*_, *RAI*, Ψ_*g50*_, Ψ_*x50*_, *k*_*xl,max*_, α, λ, *E*_0_}, represents the input parameters, and θ_0_ is the baseline parameter value. The fractional perturbation was set to_*h*_ = 0. 0005. Each parameter was perturbed independently while all other inputs were held at their original site-specific values for the corresponding thinning scenario.

As mentioned in Section 3.3.5, the growth sensitivity exponent *a* is set to 1 by default. However, this parameter could play an important role in determining leaf area growth rates, as the response of plant growth to soil water availability may vary among species. Therefore, an additional sensitivity analysis was conducted to examine how < *g* > (under unthinned conditions) changes as *a* varies from 1 to 10.

## 4. Results

### 4.1 Soil Saturation

The effect of forest thinning on soil saturation is not uniform across sites (Fig. 2, Fig. 3). For CA-TP4, FR-Pue, and FI-Hyy, reductions in *LAI* shift soil saturation distributions *p*(*s*) toward wetter conditions (Fig. 2) and increase mean soil saturation, with both the magnitude and duration of this effect increasing with thinning intensity (Fig. 3). In contrast, US-Fmf shows only a minimal increase in soil saturation under low to moderate thinning, while more intensive *LAI* reduction can initially produce a slightly negative response relative to the unthinned reference (Fig. 3).

**Figure 2.**
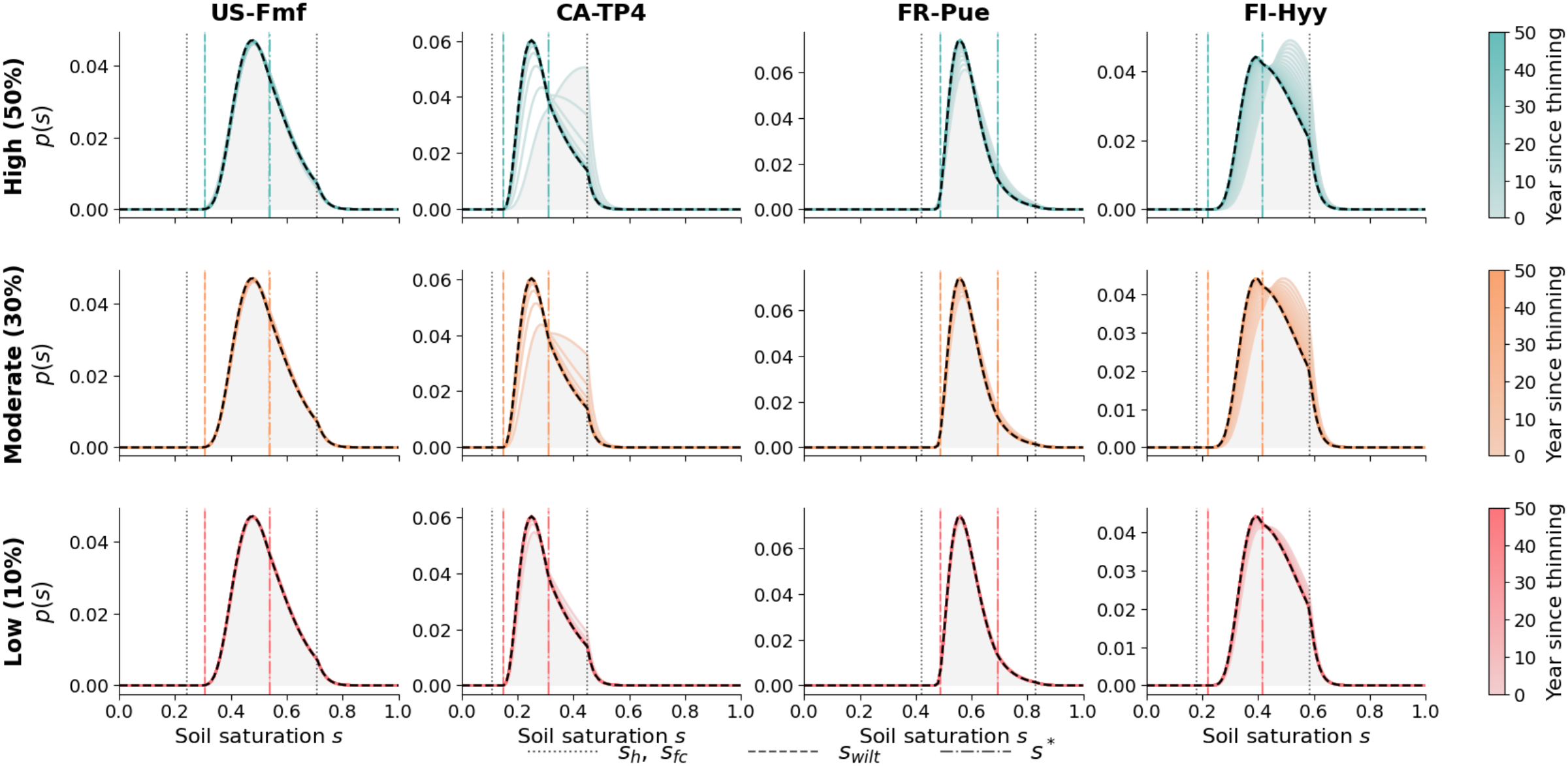
Simulated soil saturation probability density functions over 50 years. Modeled distributions of relative soil saturation (*p*(*s*)) are shown following initial leaf area index reductions of 50% (top), 30% (middle), and 10% (bottom) at four forest sites (columns). Colored curves represent successive years, while black dashed curves show the unthinned reference distributions. The shaded region represents the area under *p*(*s*) immediately after thinning. Vertical lines mark the hygroscopic threshold (*s*_*h*_), field capacity (*s*_*fc*_), wilting threshold (*s*_*wilt*_), and onset of water limitation (*s* *). Climate parameters are held constant while canopy leaf area recovers, allowing the distributions to reflect changes in vegetation structure and associated water use.

**Figure 3.**
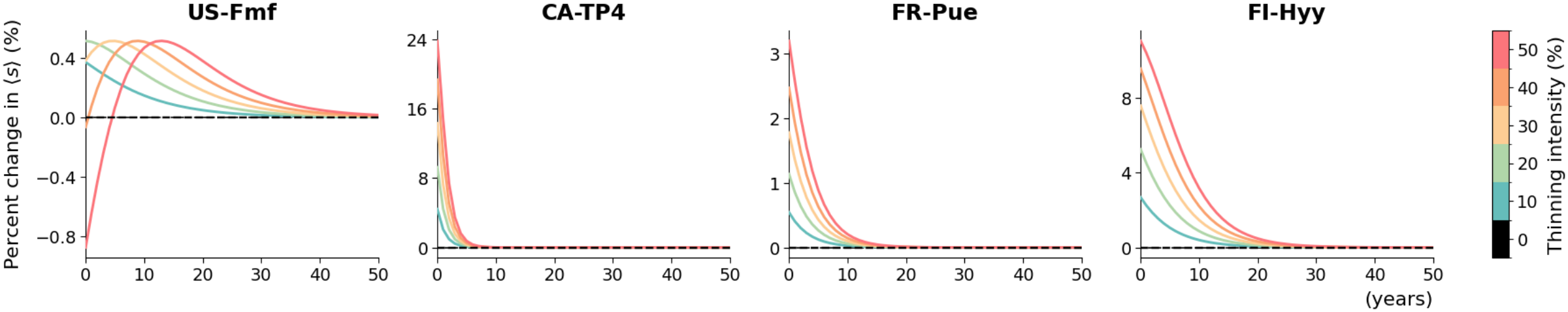
Percent changes in mean soil saturation over 50 years. Curves show percentage changes in mean relative soil saturation 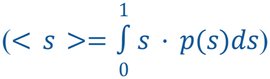, compared with the unthinned reference at four forest sites, calculated as 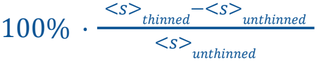.

It is important to note that the numerical value of soil saturation is strongly influenced by soil texture and water-holding capacity. For example, although CA-TP4 receives relatively high precipitation (~365 *mm growing srowiasrowin* ^−1^) it can still exhibit low soil saturation because it has sandy and well drained soils (Skubel et al., 2015). In the model, soil characteristics are represented through texture-dependent hydraulic parameters (*n, rowi*_*s*_, *s*_*h*_, *s*_*fc*_) (Section 3.3.3, A.2.1), however, the actual determinant of later water fluxes and leaf area growth is not the absolute numerical value of soil saturation, but the position of the *p*(*s*) relative to the four soil-water thresholds. For example, CA-TP4 and FI-Hyy would have the highest proportion of *E*_0_ realized because a large fraction of their *p*(*s*) lies to the right of *s* *, where transpiration reaches *T*_*ww*_ and soil evaporation continues to increase until it reaches *E*_*max*_ at *s*_*fc*_ (Fig.1.1, Fig. 2).

The effects of thinning on soil saturation are most pronounced at CA-TP4 and FI-Hyy, where a 50% reduction in *LAI* immediately shifts substantial probability mass toward the wet regime (*s* > *s* *), with the distribution peak located above *s* * (Fig. 2). Although the magnitude of the response is small, US-Fmf exhibits a hump-shaped trajectory: < *s* > initially increases as *LAI* recovers, reaches a maximum, and then declines toward the unthinned reference. This contrasts with the monotonic responses at the other sites, where lower LAI consistently corresponds to higher < *s* > (Fig. 3). This suggests that, under certain conditions, soil water retention may actually be maximized by maintaining an intermediate level of canopy cover.

### 4.2 Water Fluxes

#### 4.2.1 < *ET* >, < *E* >, **and** < *T* >

Within the model, *LAI* controls light interception, thereby influencing *E* by altering how potential evapotranspiration demand *E*_0_ is partitioned between the soil and canopy in defining maximum soil evaporation rate *E*_*max*_ (A.1.2.5). In addition to this partitioning of *E*_0_, the well water transpiration rate *T*_*ww*_ (A.1.2.5) also depends on β_*ww*_ (A.1.2.3), in which *LAI* acts as a scaling factor for its component *k*_*p,max*_ (A.1.2.1). A reduction in *LAI* therefore lowers both stand-level canopy transpiration demand and hydraulic capacity, thereby decreasing *T*_*ww*_ while increasing *E*_max_. Moreover, as thinning shifts *p*(*s*) (Section 4.1), the response of *ET* would also reflect changes in soil water supply/availability (Fig.1.2).

Across all four sites, < *E* > increases with thinning intensity (Fig. 4 (i-iv), panel a). This occurs even at US-Fmf, where *p*(*s*) shifts slightly toward lower soil saturation following intense thinning (Fig. 2), suggesting that increased radiation reaching the soil surface promotes evaporation sufficiently to outweigh the limiting effect of reduced soil water availability. In contrast, < *T* > decreases with increasing thinning intensity as the loss of leaf area reduces the total transpiring surface and thus canopy conductance, despite greater water availability at sites where soil saturation increases (Fig. 4 (i-iv), panel a, Fig. 2). Overall, the direct structural effects of *LAI* reduction generally dominate over the effects of changes in soil saturation in determining the responses of both soil evaporation and stand transpiration.

**Figure 4.** Simulated water fluxes over 50 years following forest thinning. (a) Mean evapotranspiration < *ET* >, transpiration < *T* >, and soil evaporation < *E* >. The gray dashed line indicates the observed mean growing-season *ET* from FLUXNET. (b) Relative changes in mean transpiration and soil evaporation compared with the unthinned reference, calculated as 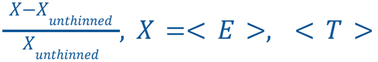.(c) Mean transpiration and soil evaporation expressed as fractions of their respective well-watered and maximum rates 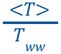 and 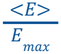, with *T*_*ww*_ and *E*_*max*_ converted to the same units as the mean fluxes. (d) Mean transpiration per unit leaf area index 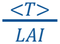. (e) Mean canopy interception < *I* >.

**Figure 4. (i).**
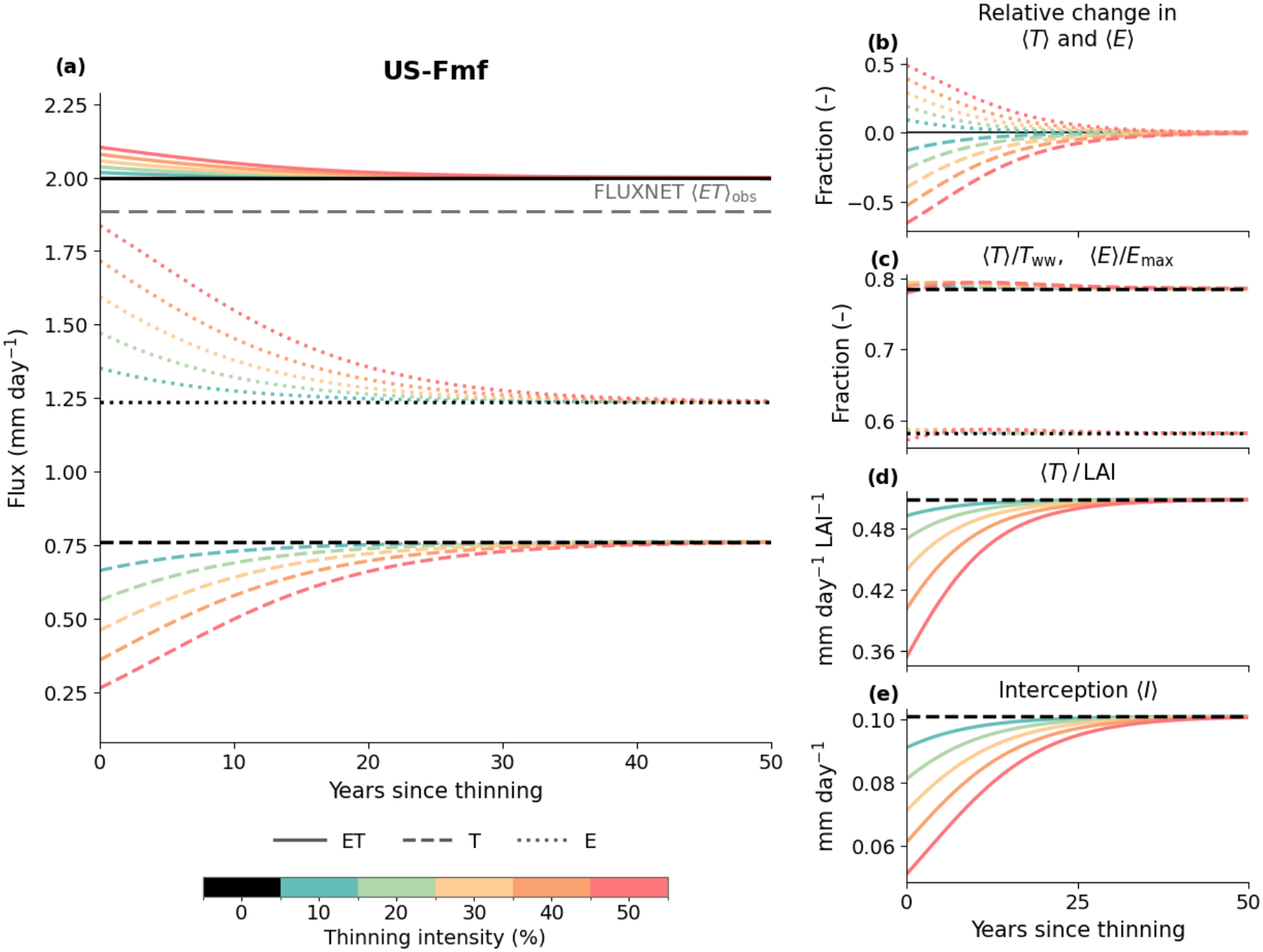
US-Fmf.

**Figure 4. (ii).**
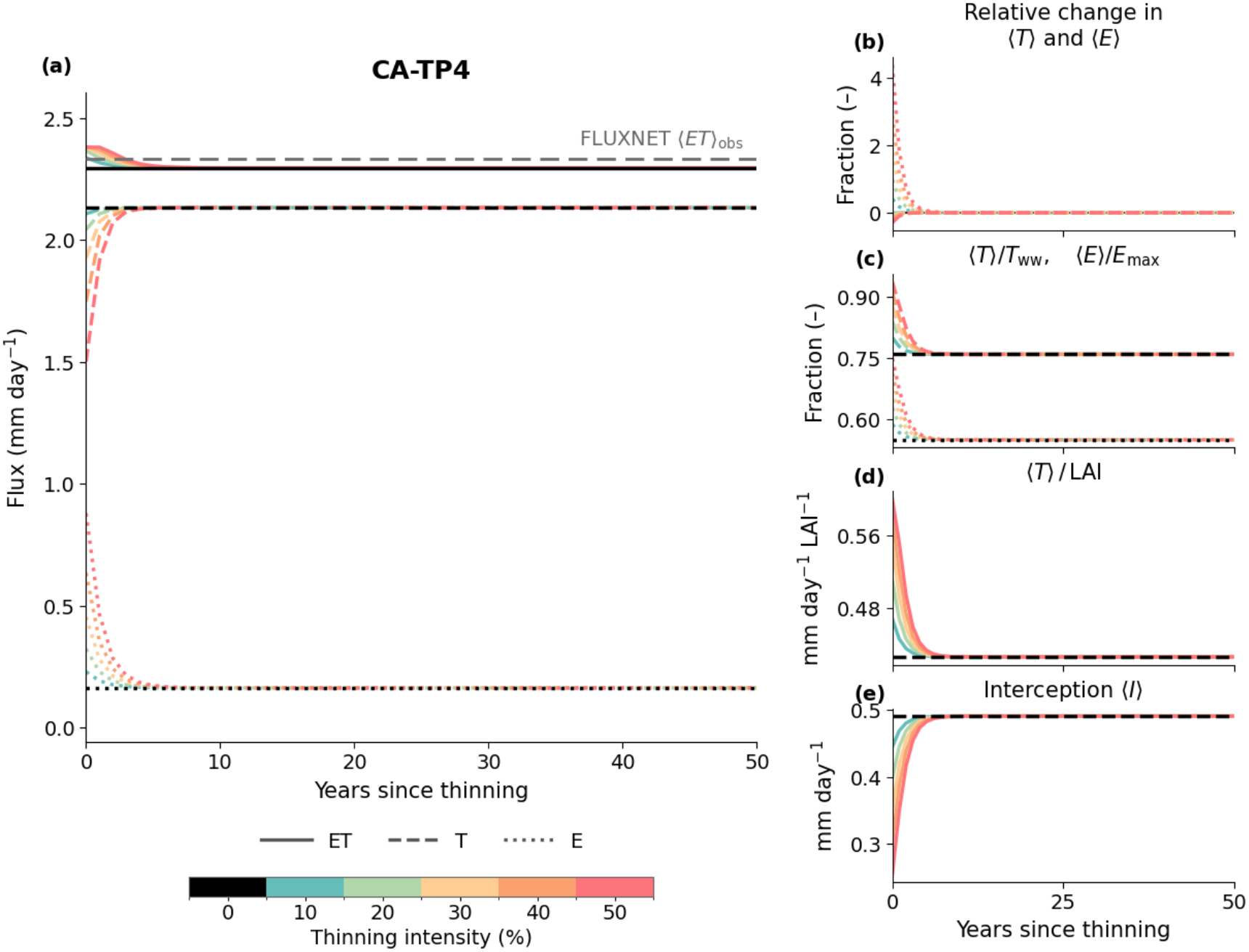
CA-TP4.

**Figure 4. (iii).**
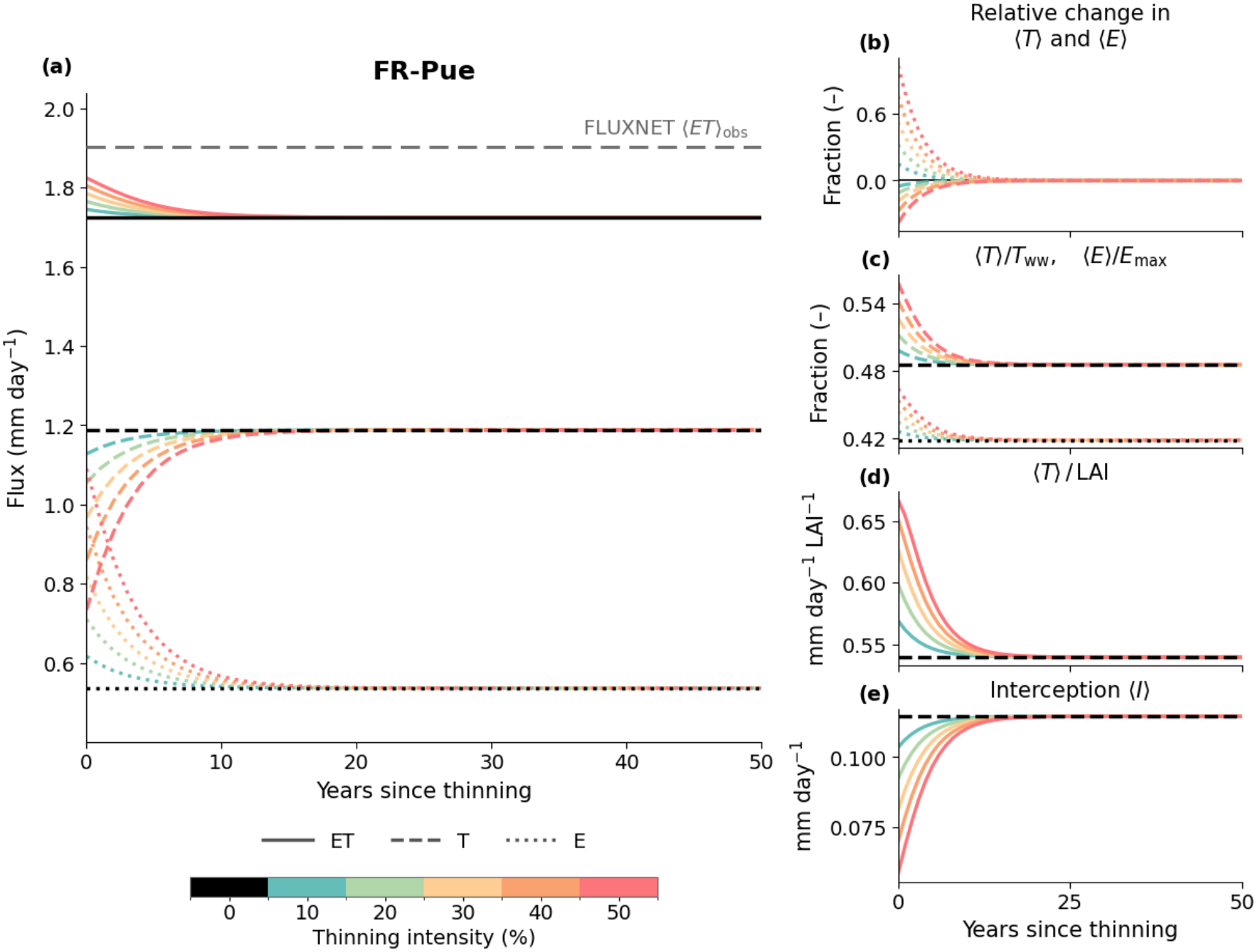
FR-Pue.

**Figure 4. (iv).**
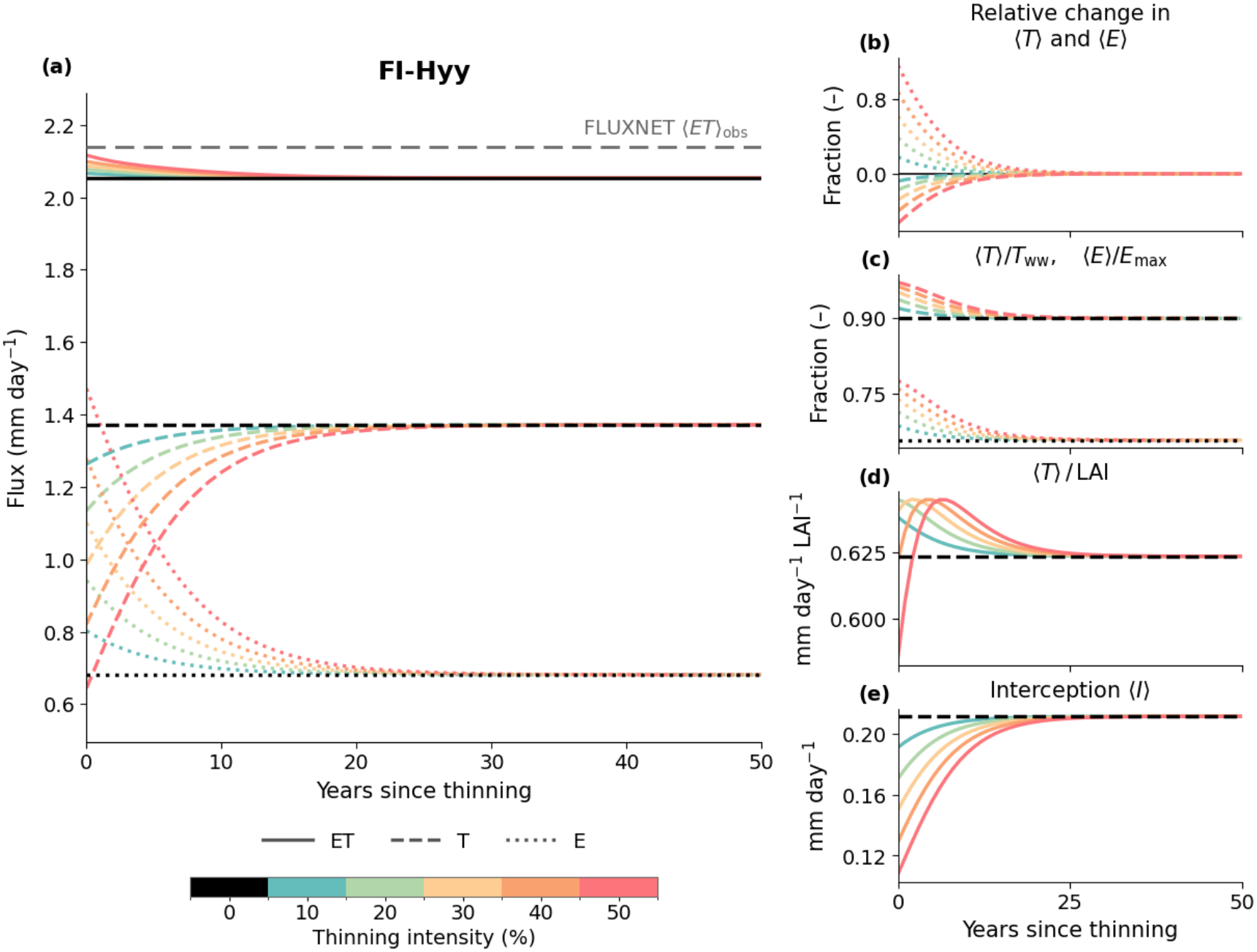
FI-Hyy.

The response of < *ET* > thus depends on the relative magnitudes of the opposing changes in soil evaporation and stand transpiration. Across all four sites, < *ET* > remains relatively stable following thinning, with slight increases at all sites, implying that the absolute increase in < *E* > slightly exceeds the absolute decrease in < *T* > (Fig. 4 (i-iv), panel a, Fig. 2).

The plots also show the observed mean growing-season *ET* from FLUXNET data. Although the simulated values differ slightly from the observations, the model broadly reproduces their magnitude despite its simplified analytical representation (Fig. 4 (i-iv), panel a).

#### 4.2.2 Sensitivity of < *T* > and < *E* > to thinning

At CA-TP4, FR-Pue, and FI-Hyy, the relative increase in < *E* > exceeds the magnitude of the relative decrease in < *T* > following thinning, indicating a greater proportional response of soil evaporation to *LAI* reduction (Fig. 4 (ii-iv), panel b). These three sites also exhibit shifts in *p*(*s*) toward wetter conditions following thinning (Fig. 2, Fig. 3), allowing a greater fraction of *E*_max_ to be realized, as reflected in 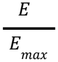 the increase in (Fig. 4 (ii-iv), panel c). Thus, improved soil water availability and increased *E*_*max*_, as discussed in section 4.2.1, act together to enhance actual soil evaporation. In contrast, although both *T*_*ww*_ and < *T* > decrease following thinning, the increase in 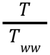 implies that despite the reduction in transpiration capacity/demand, actual transpiration decreases proportionally less due to improved soil water availability (Fig. 4 (ii-iv), panel c). This soil water compensating effect appears particularly pronounced at CA-TP4, consistent with it having the largest increase in soil saturation (Fig. 2, Fig. 4 (ii), panels b&c). At US-Fmf, the opposing relative responses of *T* and *E* are more symmetric (with *T* even showing a slightly larger proportional response) and the negligible changes in 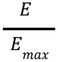 and 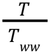 are consistent with the limited (even slightly negative) soil saturation response at this site (Fig. 4 (i), panels b & c).

Another possible contributing factor is the nonlinear, saturating relationship between *LAI* and canopy light interception. The complementary fraction reaching the soil is given by 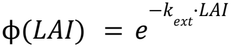. For a proportional *LAI* reduction (*rowi*), its relative increase is:

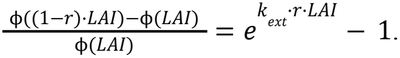

Thus, the same proportional thinning would produce a larger relative increase in soil energy at higher *LAI*_0_, although this increase is measured against a smaller baseline and does not necessarily reflect a larger absolute change in ϕ(*LAI*), whose slope is given by 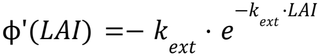. For example, with *k*_*ext*_ = 0. 55, a 50% *LAI* reduction from 1.5 to 0.75 at US-Fmf increases the soil energy fraction by approximately 22.4 percentage points, compared with 18.9 percentage points for a reduction of *LAI* from 5 to 2.5 at CA-TP4. Nevertheless, the corresponding relative increases are approximately 51% and 296%, respectively. Thus, the much larger relative evaporation response at CA-TP4 (Fig. 4 (ii), panel b) may also just reflect its smaller baseline and the model’s nonlinearity.

#### 4.2.3 Transpiration of the Remaining Stand

Here, 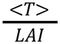 is used to characterize mean leaf level transpiration by the remaining foliage. At CA-TP4 and FR-Pue, the ratio increases following thinning, with the largest increases under the strongest treatments. Thus, although total stand < *T* > decreases, each unit of remaining leaf area transpires more water on average (Fig. 4 (ii&iii), panel d). In contrast, at US-Fmf, the ratio instead decreases, indicating that stand transpiration declines proportionally more than *LAI* (Fig. 4 (i), panel d). These contrasting responses in 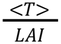 are consistent with the direction and magnitude of thinning-induced changes in soil saturation across these sites (Fig. 2) and the resulting compensatory effects on canopy transpiration, as discussed in Sections 4.2.1 and 4.2.2.

At FI-Hyy, the response depends on thinning intensity. For example, stronger thinning produces a hump-shaped trajectory, with the 50% treatment initially falling below the reference before rising to a temporary peak above it, though the ratio varies over a very narrow range (~0.06 *mm day*^−1^) (Fig. 4 (iv), panel d). One possible explanation is that actual transpiration at FI-Hyy is already close to *T*_*ww*_ even before thinning: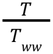 is ~ 0.9 in the unthinned stand and rises to ~0.97 under the strongest treatment. This is consistent with FI-Hyy being a relatively wet site, where a substantial fraction of the initial *p*(*s*) lies above *s* * (Fig. 2), leaving limited scope for additional soil water to further alleviate transpiration limitation. This is also reflected in the more pronounced decline in stand-level < *T* > at FI-Hyy following thinning, possibly due to the weaker compensating effect of improved *s* (Fig. 4 (iv), panel a).

#### 4.2.4 Interception

Rainfall interception decreases monotonically with increasing thinning intensity, as reduced canopy cover allows more rainfall to reach the soil as throughfall (Fig. 4 (i-iv), panel e) (Fig.1.2).

Overall, thinning has a stronger effect on the individual components and partitioning of *ET* than on its total magnitude. Although total stand-level transpiration decreases, reduced competition for water and improved soil water availability may allow the remaining foliage to partially compensate by transpiring at higher rates. At the same time, increased radiation reaching the soil surface enhances soil evaporation, offsetting the reduction in stand-level transpiration and resulting in relatively small changes in total ET.

### 4.3 *LAI* Dynamics

For all sites, the *LAI* trajectories show asymptotic recovery toward the unthinned reference, with higher thinning intensity generally resulting in larger net annual *LAI* increments but requiring longer recovery times (Fig. 5(i–iv), panels a & b). Here, two mechanisms governing the rate of canopy recovery are captured in the model: (1) reduced light interception and photosynthetic area; (2) alleviation of water-limited growth due to potentially increased soil saturation (Section 3.2, Fig.1.2). The increase in < *g* > at CA-TP4, FR-Pue, and FI-Hyy, as well as the slight decrease under heavy thinning and the following hump-shaped response at US-Fmf (Fig. 5(i-iv), panel d) are consistent with the subsequent changes in < *s* > following thinning (Fig.3). Nevertheless, the decrease of Δ*A* (A.1.2.8) at FR-Pue, US-Fmf, and FI-Hyy reflects the more dominant effect of reduced light interception on stand-level leaf area production following thinning (Fig. 5(i, iii, iv), panel c). At CA-TP4, Δ*A* initially remains above the unthinned reference under low-intensity thinning and gradually declines toward it, whereas under higher-intensity thinning, Δ*A* first increases to an early peak before returning (Fig. 5(ii), panel c). Two possible factors could contribute to this pattern: (1) Compared with the other sites, the increase in < *g* > at CA-TP4 is greater in magnitude (Fig. 5(ii), panel d), consistent with the most substantial increase in soil saturation observed at this site (Fig. 2, Fig. 3), (2) as discussed in Section 4.2.2, opposite to the response of light reaching the ground, the same percentage reduction in *LAI* results in a smaller proportional decrease in canopy light interception when the initial *LAI*_0_ is high. Thus, the negative effect of reduced light interception on Δ*A* may be less pronounced and fully offset by the increase in < *g* >.

**Figure 5.** Simulated *LAI* dynamics over 50 years following forest thinning. (a) Leaf area index (*LAI*). (b) Annual net LAI increment after accounting for leaf turnover. (c) Growing season leaf area increment (Δ*A*). (d) baseline daily leaf area growth rate (< *g* >). (e) Growing-season increment per unit leaf area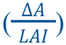.

**Figure 5. (i).**
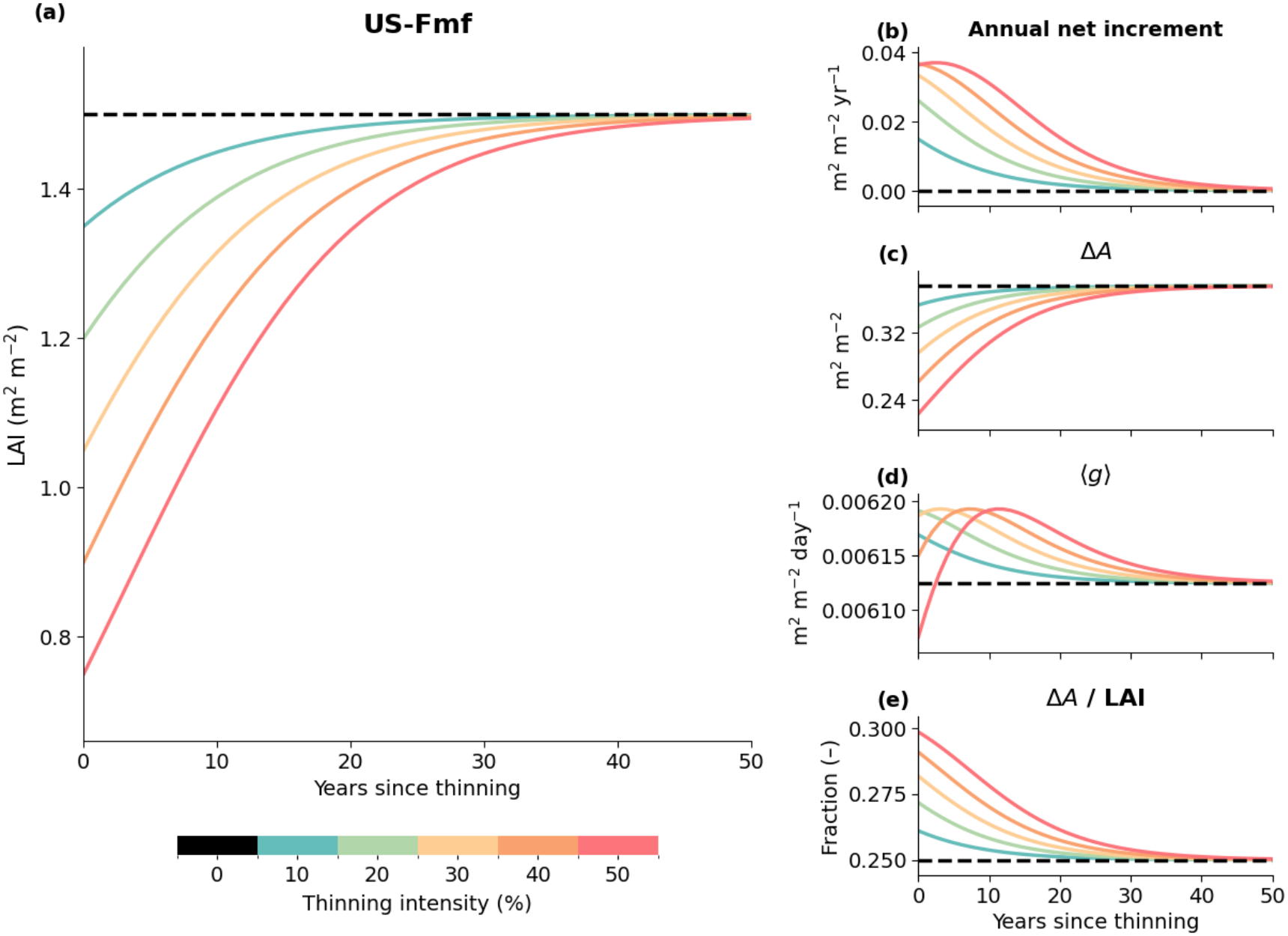
US-Fmf.

**Figure 5. (ii).**
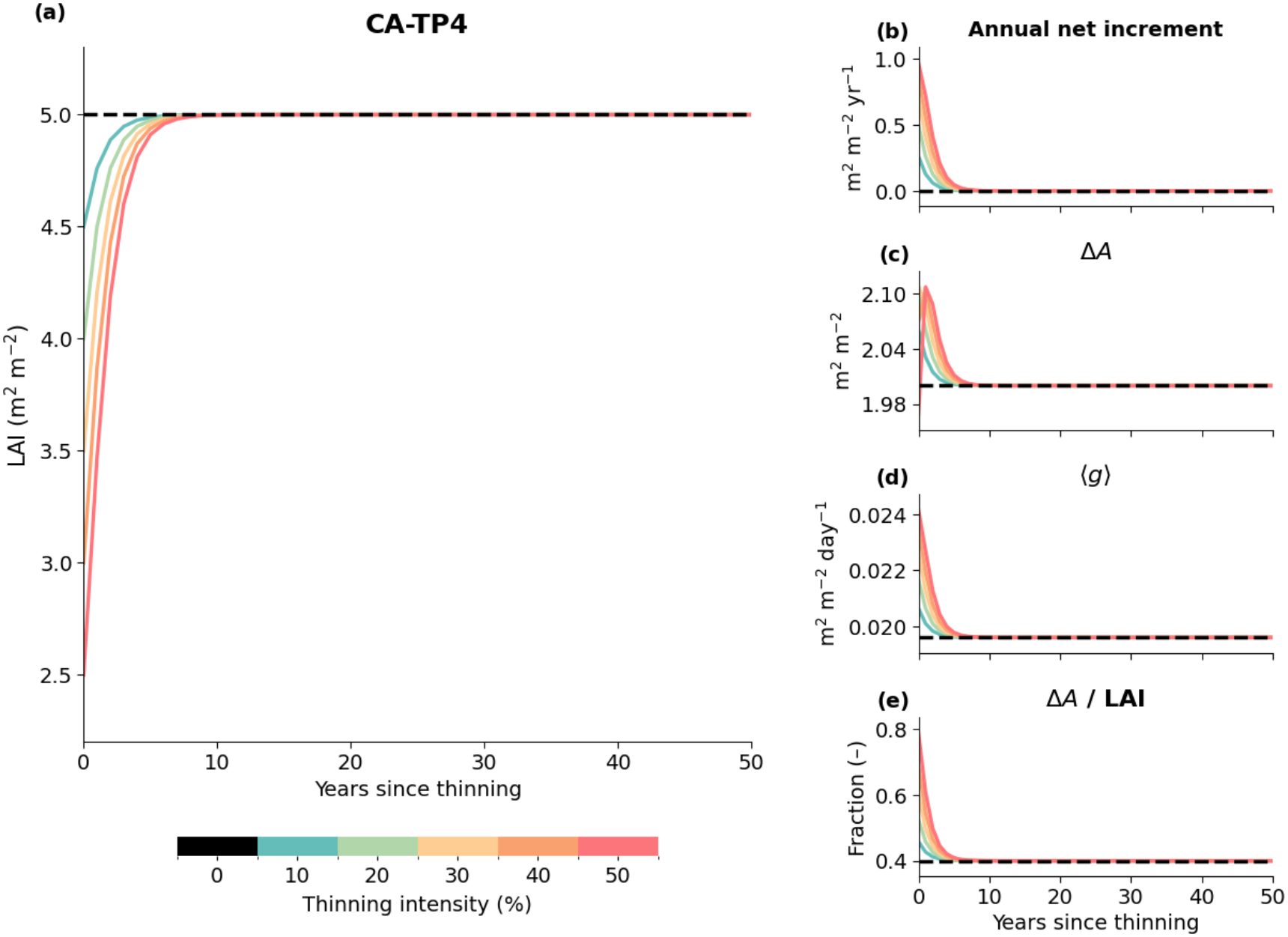
CA-TP4.

**Figure 5. (iii).**
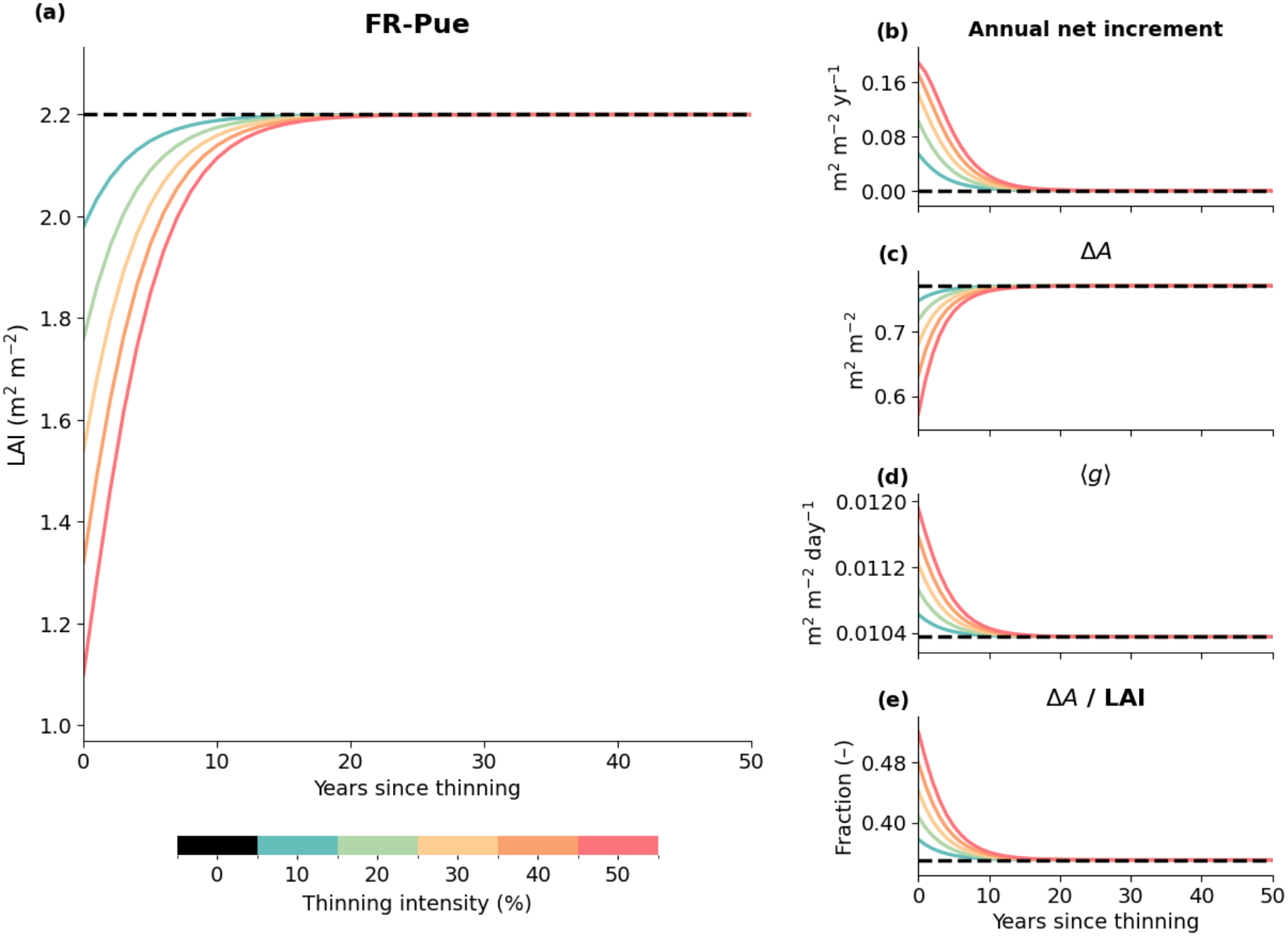
FR-Pue.

**Figure 5. (iv).**
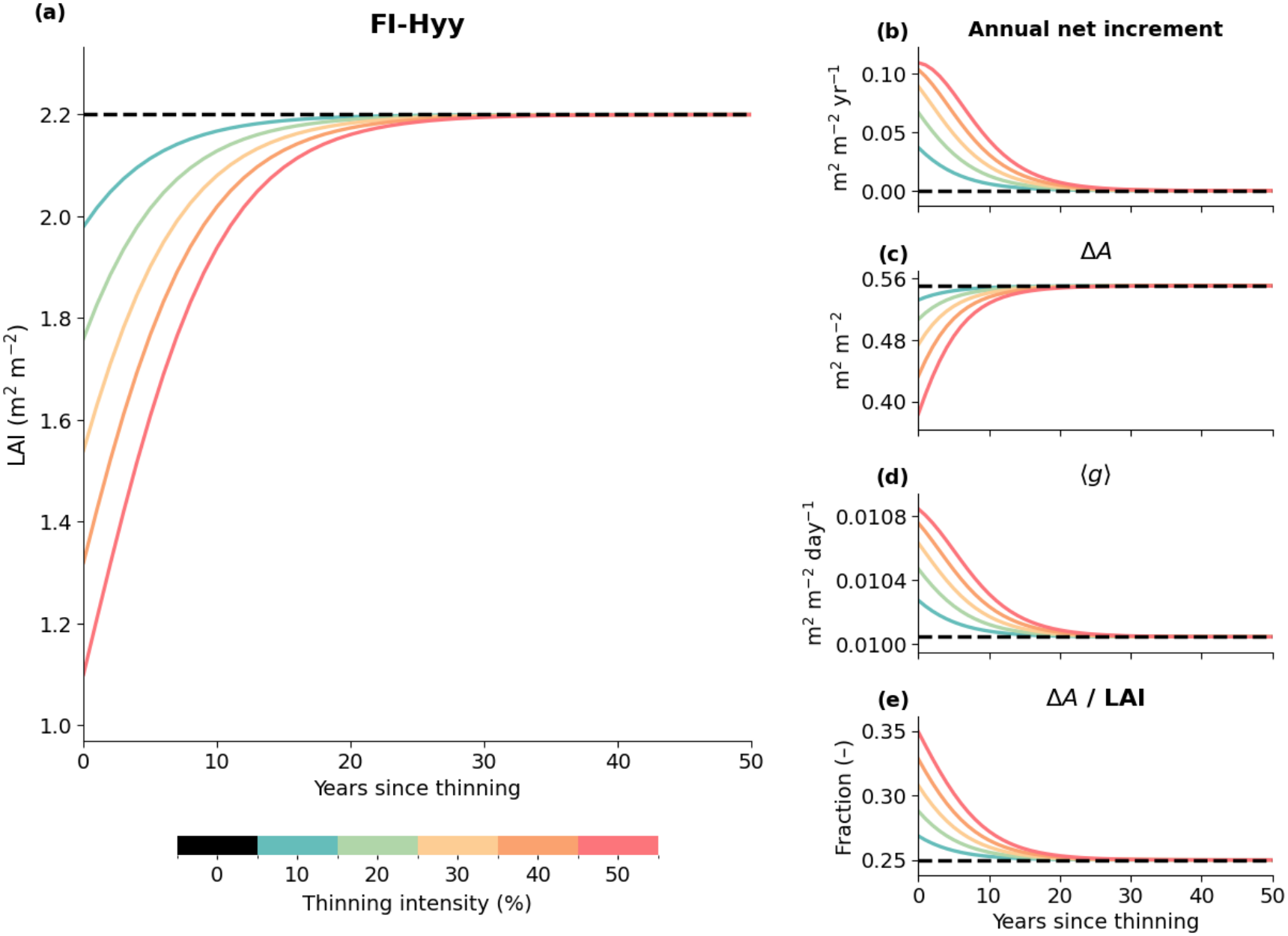
FI-Hyy.

The combined effects of a greater increase in < *g* > and a small relative reduction in canopy light interception help explain the exceptionally rapid canopy recovery at CA-TP4 (Fig. 5(ii), panel a), also with more leaf area remaining after thinning, CA-TP4 would naturally have a higher stand-level growth rate. Another contributing factor is that CA-TP4 also has a higher *g*_+_ (Table.1). *g*_+_ is calibrated numerically by requiring annual leaf-area production to perfectly balance leaf shedding in the unthinned reference (Section 3.3.5). Consequently, its value depends on the prescribed turnover rate β: with all other conditions held constant, a higher β requires a larger *g*_+_ to maintain the same equilibrium *LAI*. This balance is consistent with leaf economics theory, whereby shorter-lived foliage requires greater leaf production for replacement and is generally associated with higher specific leaf area (*rowiLA*) and thus lower biomass investment per unit leaf area (Gibert et al., 2016). The dominant species at CA-TP4, eastern white pine (*Pinus strobus*), has a reported *rowiLA* of approximately 113 *rowim*^2^ *g* ^−1^ (Wang & Chen, 2012), considerably higher than the average value for *Pinus* species (Berner & Law, 2016). Thus, although through a very limited representation, the model is able to capture some aspects of plant growth strategies. Also, growing-season length is another contributing factor.

Despite some decreases in Δ*A*, all sites show an increase in 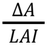 following thinning, implying that each remaining unit of *LAI* supports more leaf growth (Fig.5 (i-iv), panel e).

**Table 1.**
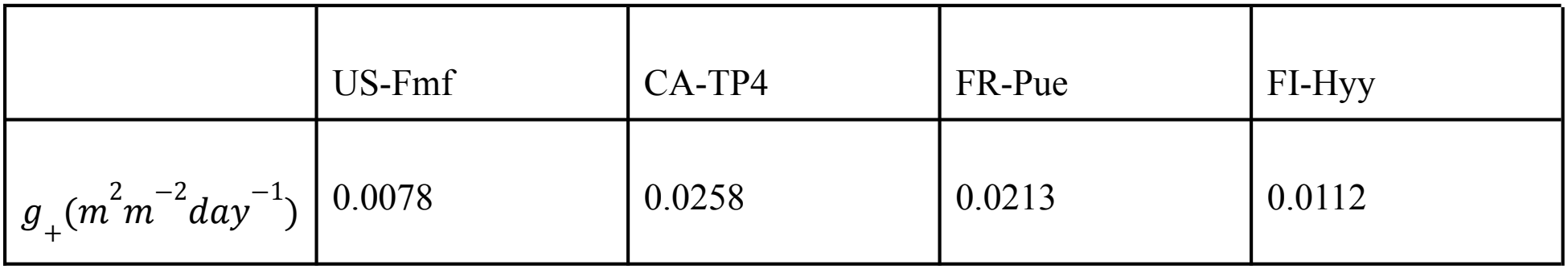
Maximum leaf area growth rate (*g*_+_) Values represent the maximum leaf area growth rates *g*_+_ (*m*^2^ *m*^−2^ *day*^−1^) at each site, numerically calculated following the method described in Section 3.3.5.

|  | US-Fmf | CA-TP4 | FR-Pue | FI-Hyy |
|---|---|---|---|---|
| *g*<sub>+</sub>(*m*<sup>2</sup>*m*<sup>−2</sup>*day*<sup>−1</sup>) | 0.0078 | 0.0258 | 0.0213 | 0.0112 |

### 4.4 Sensitivity Analysis

#### 4.4.1 Soil and roots (*Z* & *RAI*)

Key outputs (< *ET* >, < *T* >, < *E* >, < *s* >) responded minimally to changes in effective rooting-zone depth *Z*_*m*_ and *RAI* (Fig. 6 (i-iv), panels a-d). A larger *Z*_*m*_ increases soil-water storage capacity, so the same amount of water gained through infiltration or lost through *ET*/drainage produces a smaller change in soil saturation. *Z*_*m*_ therefore alters the magnitude and timing of soil-moisture fluctuations without directly changing rainfall input or atmospheric demand. This interpretation is consistent with the model simulations by Laio et al. (2001), in which rooting depths of 0.3 and 0.9 m yielded approximately the same < *s* > but different *p*(*s*), with shallower soils exhibiting greater temporal fluctuations in soil saturation and broader *p*(*s*) distributions.

**Figure 6.**
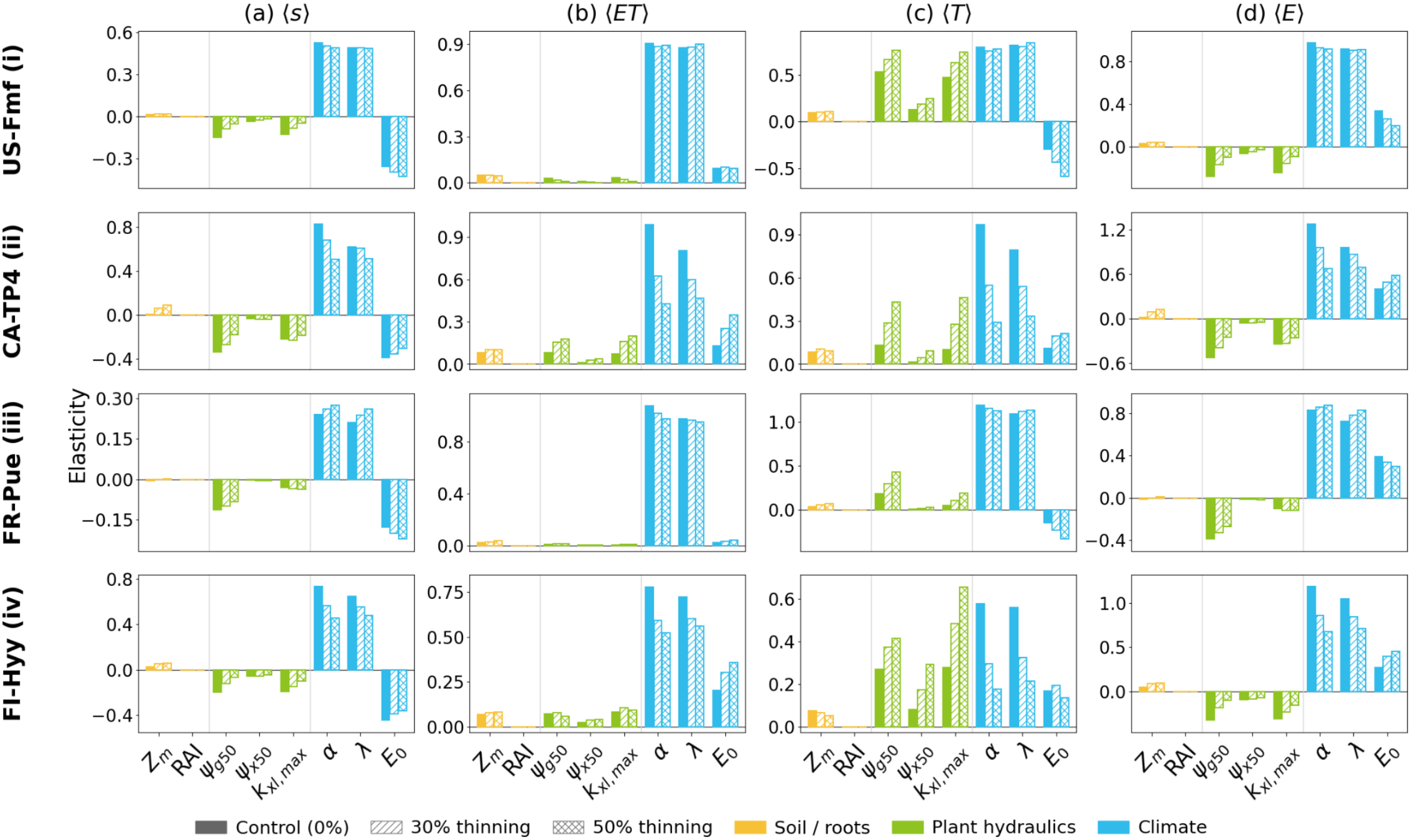
Elasticity analysis. The y-axis represents the elasticity of first-year < *s* > (a), < *ET* > (b), < *T* > (c), and < *E* > (d) with respect to the input parameters (x-axis), grouped into soil and root properties, plant hydraulic traits, and climatic inputs. Elasticity is calculated separately under 0%, 30%, and 50% thinning scenarios.

*RAI* enters the model through *rowi*_*srowi,max*_ (A.1.2.1). However, transpiration in the model is generally more strongly constrained by plant xylem conductance and stomatal regulation, and soil-to-root conductance has relatively little influence on plant water uptake capacity (Bassiouni et al., 2023).

#### 4.4.2 Plant hydraulics (Ψ_*g50*_, Ψ_*x50*_, *k*_*xl,max*_)

More negative Ψ_*g50*_ and Ψ_*x50*_, together with greater *k*_*xl,max*_, generally increase plant water uptake capacity by allowing stomatal and xylem conductance to be maintained at more negative water potentials and enhancing maximum xylem hydraulic conductivity, respectively, and thus transpiration < *T* >. In general, < *T* > is more sensitive to Ψ_*g50*_ than to Ψ_*x50*_. This is consistent with the mechanisms described by Bassiouni et al. (2023). Specifically, Ψ_*g50*_ influences all four nondimensional Π groups governing plant water uptake (A.1.2.2), including plant water flux control Π_*rowi*_, which was identified as the dominant factor influencing plant water uptake strategies. In contrast, Ψ_*x50*_ affects only hydraulic risk tolerance Π_*R*_, whose influence is generally smaller and only becomes more pronounced when plant water transport capacity is strongly limiting relative to atmospheric demand (Bassiouni et al., 2023). Similarly, *k*_*xl,max*_ determines *rowi*_*p,max*_ (A.1.2.1) and thus influences the capacity of plants to transport water from roots to the canopy.

Across all sites, the elasticity of < *T* > to hydraulic traits generally increases with thinning (Fig. 6 (i-iv), panel c). One plausible explanation is that thinning may alleviate soil water limitation by shifting *p*(*s*) toward wetter states (Fig. 2), where a greater proportion of *T*_*ww*_ is realized, allowing these hydraulic traits that determine β_*ww*_ (A.1.2.3) and thus *T*_*ww*_ (A.1.2.5) to exert a stronger influence on < *T* >.

However, the degree of elasticity amplification depends on site-specific *p*(*s*) feedback and the nonlinear response of transpiration to soil saturation. Lower baseline transpiration following thinning may also contribute to larger relative changes without necessarily indicating larger absolute responses.

The response of < *E* > to changes in plant hydraulic traits is mediated by changes in soil saturation, and generally opposes the response of < *T* >, as greater plant water uptake reduces soil-water availability and suppresses evaporation. These opposing changes largely offset one another, resulting in the relatively weak sensitivity of < *ET* >. Thus, at a fixed *LAI*, plant hydraulic traits would primarily influence the partitioning of *ET* between transpiration and evaporation (Fig. 6 (i-iv), panels a, b, and d).

### 4.4.3 Climatic inputs (α, λ, *E*_0_)

Precipitation inputs α and λ directly affect soil-water recharge and, consequently, soil saturation and water fluxes. In general, model outputs exhibit slightly greater elasticity with respect to α than to λ (Fig. 6 (i-iv)). This difference may partly result from the canopy interception mechanism, which is represented as a fixed depth of rainfall intercepted per event (_*h rowi*_, A.1.1.4). Larger rainfall events therefore have a smaller intercepted fraction, whereas increasing λ increases the number of rainfall events without changing the fraction of rainfall intercepted at a fixed *LAI* and α. The corresponding positive responses of < *ET* >, < *E* >, and < *T* > are therefore mediated by increased soil saturation. Increasing *E*_0_ raises potential water loss, thereby accelerating soil drying.

The effects of thinning on the elasticities of selected model outputs with respect to climatic inputs vary across sites, reflecting differences in soil water availability and the relative importance of water-supply versus atmospheric-demand limitations. At CA-TP4 and FI-Hyy, all examined outputs exhibit substantial reductions in elasticity to α and λ following thinning, with < *T* > showing the most pronounced changes (Fig. 6 (ii & iv)). One plausible explanation is that, at these two sites, a large proportion of *p*(*s*) is initially concentrated in the wetter soil-saturation range (Fig. 2), and thinning also results in relatively larger increases in soil saturation (Fig. 3), further alleviating soil-water limitations. As a greater proportion of atmospheric evapotranspirative demand is already met, further increases in soil-water recharge may produce smaller proportional increases in *ET*. This interpretation is consistent with the higher positive elasticity of < *ET* > to *E*_0_, following thinning (Fig. 6 (ii & iv), panel b), suggesting that atmospheric evaporative demand becomes a more important constraint on *ET* at these two sites. In contrast, at FR-Pue and US-Fmf, elasticity to precipitation parameters varies relatively little across thinning conditions, consistent with the relatively small or even negative changes in soil saturation following thinning (Fig. 3, Fig. 6 (i & iii)). At these sites, transpiration may remain more strongly constrained by soil-water availability before being limited by atmospheric evapotranspirative demand, as reflected in the weak response of < *ET* > to *E*_0_ (Fig. 6 (i & iii), panel b). Increasing *E*_0_ raises *T*_*ww*_ (A.1.2.5) but also accelerates soil-water depletion, potentially reducing actual plant water uptake through greater soil-water stress. This may explain the negative elasticity of < *T* > to *E*_0_ at these sites (Fig. 6 (I & iii), panel c).

#### 4.4.4 Growth sensitivity exponent (*a*)

< *g* > decreases with increasing *a* due to the stronger suppression of growth under water-limited conditions at higher exponents (Fig. 7). This can have a significant effect on growth, especially at sites such as FR-Pue, where a large proportion of the soil-moisture distribution *p*(*s*) lies within the water-limited range (Fig. 2). Therefore, the choice of *a* can affect model behavior, and incorporating species-specific measurements or empirically derived relationships between growth and soil moisture could improve model predictions.

**Figure 7.**
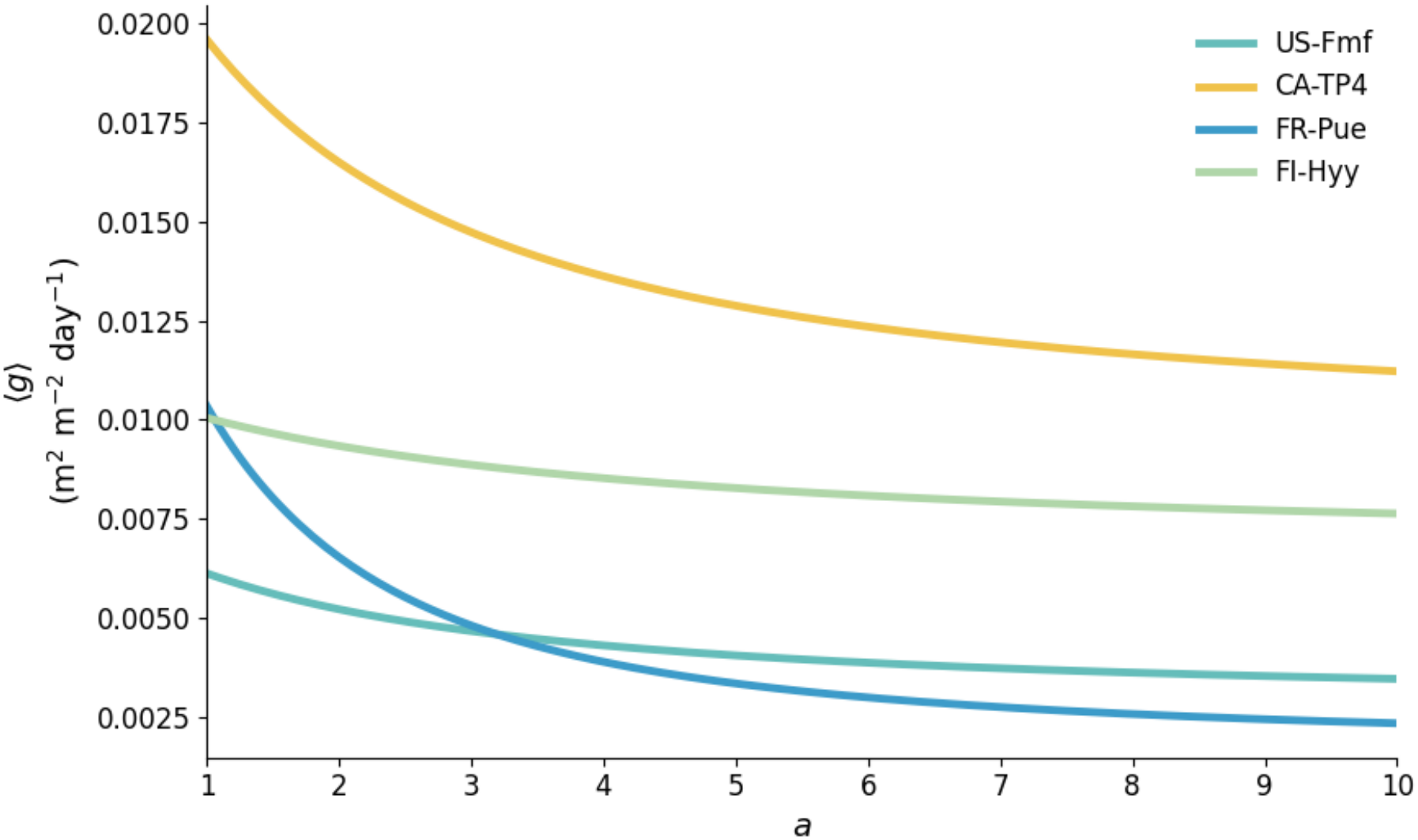
Sensitivity of mean growth rate to the growth sensitivity exponent *a*. Curves show < *g* > as a function of the growth sensitivity exponent *a*, ranging from 1 to 10, across four forest sites under unthinned reference conditions.

## 5. Discussion

### 5.1 Soil Saturation Response to Thinning

Although this study examines only four sites, the contrasting soil saturation responses highlight the importance of local climatic, hydrological, and forest structural characteristics in determining the water conservation benefits of forest thinning and the effects of thinning intensity. Under certain conditions, more intensive thinning does not necessarily conserve more soil water and may even have negative effects. Similar variability in soil saturation responses to thinning has also been widely documented in previous studies. In a global meta-analysis of soil-moisture responses across thinning intensities of 1–46% stand-density removal, del Campo et al. (2022) reported an average 14% increase in soil moisture following thinning. Similarly, Bian et al. (2026) found mean increases of 9% in planted coniferous forests and 23% in natural coniferous forests across various thinning intensities. Another meta-analysis of coniferous, broadleaf, and mixed forest plantations reported approximate mean increases in soil moisture of 3%, 13%, and 9% under thinning intensities of 0–30%, 30–60%, and 60–100%, respectively (Zhang et al., 2023).

However, all three meta-analyses reported substantial variation in soil-moisture responses depending on thinning treatment, forest type, and the depth at which soil moisture was measured, including cases where thinning produced negligible or even negative effects (del Campo et al., 2022; Bian et al., 2026; Zhang et al., 2023). del Campo et al. (2022), for example, found that thinning effects were moderate or limited in drier climates, where transpiration is already strongly suppressed by soil-water deficits, masking the additional water savings produced by thinning. In addition, in some settings, low precipitation may limit the extent to which increased throughfall following thinning translates into greater soil-water storage (Bian et al., 2026). This may help explain the relatively small soil saturation response at FR-Pue (Fig. 2, Fig. 3), which experiences a dry mediterranean growing season with average precipitation of approximately 0.002 m per day. In comparison, FI-Hyy receives approximately 0.0025 m of precipitation per day during the growing season. This is also consistent with the model’s formulation of canopy interception as a fixed depth per rainfall event proportional to *LAI* (A.1.2.4). Thus, despite having the same *LAI*_0_, FI-Hyy exhibits a larger reduction in < *I* > than FR-Pue under the same thinning intensity (Fig. 4 (iii–iv), panel e).

Bian et al. (2026) also reported decreases in soil moisture following thinning in three studies involving forests with relatively low initial stand densities. The authors suggested that further increases in canopy openness and reduced shading could increase the energy reaching the soil surface, thereby enhancing soil evaporation and reducing soil moisture (consistent with the illustration in Fig.1.2) (Bian et al., 2026).

This mechanism is consistent with the small or even slightly negative soil saturation response at US-Fmf, which has a low *LAI*_0_ of 1.5 and where soil evaporation accounts for a dominant proportion of total *ET* (Fig. 2, Fig. 3, Fig. 4 (i)).

When evaluating the potential water conservation benefits of forest thinning, its effects under climatic variability are an important factor to consider. For example, Tatum et al. (2025) found that mean soil water availability in thinned stands at the Fort Valley Experimental Forest in northern Arizona remained relatively stable despite substantial interannual precipitation variability. They suggest that reduced competition for soil water and lower stand-level transpiration demand following thinning may allow soil water to be depleted more slowly during dry years. Another study in Lassen National Forest, California, found that thinning helped maintain higher soil-water storage and delayed soil-moisture depletion, particularly during dry years (Hardage et al., 2022). The simulations assume stationary interannual climatic conditions; nevertheless, the elasticity analysis indicates that thinning could reduce the sensitivity of soil saturation to precipitation perturbations at some sites (CA-TP4 & FI-Hyy, Fig. 6 (ii & iv)). This may suggest that thinning could help buffer the effects of rainfall variability on soil water storage.

However, without a direct comparison of thinning responses under paired dry and wet conditions or simulations incorporating interannual climatic variability, this hypothesis remains uncertain.

Because this study includes simulations of only four forest sites, the sample size is too limited to draw general conclusions about between-site patterns in soil-saturation responses to thinning. Nevertheless, the results suggest that these responses are not necessarily monotonic with thinning intensity and may depend strongly on forest structure and climatic conditions.

### 5.2 *ET* Response to Thinning

In general, soil evaporation increases while canopy transpiration decreases following thinning (following the mechanism in Fig.1.2), with these two opposing changes largely offsetting one another, resulting in relatively stable total *ET* (Section 4.2). This is broadly consistent with the findings and proposed compensatory mechanisms of del Campo et al. (2022), who reported that total *ET* was approximately 4% higher in unthinned control stands than in thinned stands, a difference that is not statistically significant or conclusive (del Campo et al., 2022). As thinning may alleviate soil water limitation, the remaining foliage may transpire at a higher rate, partially offsetting the reduction in stand-level transpiration, as reflected by the increase in 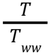 and 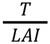(Sections 4.2.2 and 4.2.3). Increased tree-level transpiration following thinning has also been documented in several field studies (Aslan et al., 2024; Simonin et al., 2007; Lagergren et al., 2008; Gebhardt et al., 2014).

Liu et al. (2018), studying a managed loblolly pine plantation in the coastal plain of North Carolina, also found relatively small changes in total *ET* following thinning. The authors attributed this limited response partly to the forest being predominantly energy-limited rather than water-limited, as shallow groundwater provided an additional water source and net radiation strongly influenced *ET* variability.

This finding may provide a possible connection to the sensitivity analysis in Section 4.4.3, where the two relatively wetter sites, CA-TP4 and FI-Hyy, exhibit greater sensitivity of *ET* to *E*_0_ than the other two sites, particularly under thinning (Fig. 6, panel b). Both the Turkey Point region and Hyytiälä have been characterized as relatively energy-limited environments, with Hyytiälä in particular described as water-demand limited (which is also reflected in its relatively low *E*_0_, A.2) (Arango Ruda & Arain, 2025; Nadal-Sala et al., 2021). This pattern suggests that *E*_0_ may play a more important role in regulating total *ET* where water availability is less limiting. One possible explanation for why this effect is even more pronounced following thinning at these two sites is that thinning further alleviates water limitation, shifting *ET* toward even more energy-limited conditions. However, this connection remains tentative.

In addition to its weak response to changes in leaf area, total *ET* also exhibits low sensitivity to plant hydraulic traits, despite their substantial effects on transpiration (Section 4.4.2, Fig. 6, panels b, c, & d). This suggests that plant water-use traits, and perhaps, by a more speculative extension, plant functional types, may have a relatively limited influence on total ecosystem *ET*, as soil evaporation tends to change in the opposite direction to transpiration (Fig. 6, panel d). A synthesis of 167 FLUXNET sites found that climatic water supply and evaporative demand explained most of the across-site variation in annual evaporative index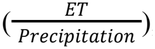, while vegetation type alone had no statistically significant additional effect (Williams et al., 2012), consistent with the substantially greater elasticity of modeled < *ET* > to climatic inputs in Fig. 6, panel b. Similarly, although conducted in a shrub–grass ecosystem rather than a forest, Wang et al. (2018) found, using a multisource surface energy-balance model, that changes in shrub–grass composition produced larger shifts in *ET* partitioning but had relatively little effect on total *ET*. They attributed the different *ET* partitionings to changes in vegetation cover and *LAI*, as well as contrasting soil water uptake strategies and transpiration patterns between shrubs and grasses, mechanisms consistent with those identified in this study (Wang et al., 2018). Similarly, Huxman et al. (2005), using a water-balance framework, suggested that vegetation changes may have relatively small effects on total *ET* especially in arid and semiarid ecosystems because most available precipitation is eventually lost through either evaporation or transpiration, though the relative contributions of *E* and *T* change substantially. This is consistent with the particularly weak sensitivity of < *ET* > to plant hydraulic traits at US-Fmf and FR-Pue, the two drier sites (Fig. 6 (i & iii), panel b).

Thus, one broader hypothesis emerging from this study is that changes in vegetation characteristics, particularly *LAI* and potentially plant hydraulic traits, may redistribute water losses among different pathways, including canopy interception, soil evaporation, and transpiration, as discussed by Huxman et al. (2005). However, the net response of total ET depends more strongly on precipitation inputs and atmospheric evaporative demand.

### 5.3 Post Thinning Canopy Recovery

*LAI* recovery in this model reflects two layers of control: soil-water availability and canopy light interception, along with site-specific growing conditions, such as growing-season length and leaf turnover rate (Section 3.1.4, Section 3.2). Therefore, it should be interpreted primarily as a mechanism for updating *LAI* within a consistent soil-saturation framework to evaluate how thinning-induced ecohydrological changes diminish over time, rather than as a direct measure of forest productivity. Although *LAI* can serve as an indirect proxy for productivity, this model does not explicitly account for species-specific photosynthetic efficiency, carbon allocation, or growth limitations imposed by nutrient availability and other environmental factors (Castro-Izaguirre et al., 2016). Furthermore, in forest management, productivity is often evaluated in terms of wood production, for example, through increased radial growth of remaining trees following thinning documented in several studies (Sohn et al., 2016). The present model is limited in its ability to capture such responses. However, this model still captures some key mechanisms through which light interception and soil-water availability control leaf-area growth. In general, greater *LAI* provides more photosynthetic surface area and increases light interception, as reflected in the overall reduction in Δ*A* following thinning (Fig. 5 (i, iii, & iv), panel c). At the same time, greater *LAI* increases self-shading, which can reduce growth efficiency per unit leaf area (Waring, 1983). This is conceptually consistent with the increase in 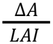 (Fig. 5 (i-iv), panel e). The model further incorporates water limitation, such that increased soil saturation following thinning can also support compensatory leaf-area growth in the remaining canopy (Fig.1.2). The relationship between *LAI* reduction and leaf-area growth is nonlinear, and the same proportional reduction in *LAI* does not necessarily produce the same change in growth across sites, as discussed in Section 4.3. This is consistent with the findings of Bouriaud et al. (2025), who showed that stand-level gross primary productivity tends to saturate once *LAI* reaches approximately 3–4.5, beyond which additional foliage contributes relatively little to stand-level photosynthesis. This may also help explain the growth response at CA-TP4, where the relatively high *LAI*_0_ limits the reduction in canopy light interception following thinning. Considering the model’s incorporation of soil-saturation controls on growth, the relatively greater increase in soil saturation following thinning at CA-TP4 completely offset the small reduction in canopy light interception by alleviating water limitations on growth (Fig. 5 (ii)).

This suggests that there might exist an intermediate range of *LAI* that balances sufficient light interception with reduced competition for soil water, thereby optimizing canopy growth. This is consistent with the conclusion of another modeling study, which suggests that vegetation canopy development is constrained by a trade-off between maximizing light absorption and avoiding excessive soil-water depletion (Kergoat, 1998).

### 5.4 Model validation against field observations

#### 5.4.1 Soil saturation

There are some discrepancies between field observations and model simulations in soil moisture responses to thinning. At US-Fmf, field measurements showed higher dry-season soil water content at both 10 cm and 50 cm depths in thinned stands than in the unthinned control (Dore et al., 2012), whereas the model predicted only minimal or slightly negative changes (Fig. 2, Fig. 3). At FI-Hyy, field observations showed an immediate increase in volumetric soil moisture across all measured soil layers following thinning. At a depth of 0.3–0.4 m, the difference was approximately 17 percentage points in volumetric soil moisture (~85% relative increase) following the ~40% thinning treatment (de Quesada, 2026), considerably larger than the approximately 9% relative increase in mean soil saturation produced by the corresponding simulation (Fig. 3). Although these quantities are not directly comparable, as they represent soil water content using different measures, the differences may still suggest that additional processes not represented in the model may influence post-thinning soil-water dynamics

#### 5.4.2 ET

In this study, across all four simulated sites, the increase in soil evaporation slightly exceeds the reduction in canopy transpiration, resulting in a small increase in total *ET* following thinning (Fig. 4, panel a). However, field observations show somewhat different responses. At the Flagstaff ponderosa pine forest (US-Fmf), *ET* generally decreased following thinning relative to the unthinned control, with a few monthly exceptions, although this difference diminished and became negligible in approximately four years (Dore et al., 2010, 2012). Similarly, at the Hyytiälä SMEAR II boreal forest (FI-Hyy), ecosystem *ET* decreased slightly following thinning, although the overall change was small (Aslan et al., 2024). In contrast, at the Turkey Point 1939 eastern white pine plantation (CA-TP4), *ET* measured after thinning was slightly higher than in the preceding years (Skubel et al., 2015). However, the ecosystem *ET* measurements at CA-TP4 and FI-Hyy were made at the same site before and after treatment without a directly comparable contemporaneous unthinned control. Therefore, the influence of interannual climatic variability cannot be entirely excluded, and the observed differences cannot be attributed conclusively to thinning.

#### 5.4.3 Leaf area growth

As discussed in Section 5.3, the representation of leaf growth in the present model is particularly simplified. Nevertheless, the simulated *LAI* recovery trajectories show some agreement with field observations. For example, at FR-Pue, field measurements conducted approximately 30 years after thinning showed little difference in *LAI* among treatments, including the unthinned control (Cabon et al., 2018). In these simulations, *LAI* approaches its pre-thinning level within approximately 10–15 years (Fig. 5 (iii), panel a), which is within the time of eventual canopy recovery, although the field measurements do not establish the actual time required for recovery. Similarly, at US-Fmf, tree *LAI* decreased from 1.5 to 0.9 immediately following thinning and recovered only slightly to approximately 1.0 four years later (Dore et al., 2010, 2012). This slow initial recovery is broadly consistent with the simulated trajectory under the 40% thinning scenario (Fig. 5 (i), panel a).

### 5.5 Study limitations

#### 5.5.1 Model Structure and Assumptions

In the present model, forest structure is simplified to a single stand-level variable, *LAI*, and *ET* is represented by soil evaporation and transpiration from the monospecific tree canopy. However, thinning is often followed by rapid growth of understory vegetation, including shrubs, herbs, and grasses, which can contribute to ecosystem *ET* and also compensate for reduced overstory transpiration (Simonin et al., 2007; Liu et al., 2018; del Campo et al., 2022; Gebhardt et al., 2014; Kassuelke et al., 2022). Understory transpiration is not explicitly accounted for in the present model, limiting its ability to capture the redistribution of water use among different vegetation layers following thinning.

Another limitation is the simplified representation of change of plant hydraulic structure following thinning. Reductions in stand-level *LAI* are not necessarily proportional to reductions in conducting sapwood area, as the relationship between leaf area and sapwood area varies among species, tree size, tree age, and stand density (Kaufmann & Troendle, 1981; Medhurst & Beadle, 2002). For example, Medhurst & Beadle (2002) found a nonlinear relationship in which larger trees supported more leaf area per unit sapwood area. Furthermore, several thinning studies have documented increases in the leaf area to sapwood area ratio with decreasing stand density (McDowell et al., 2006; Giuggiola et al., 2013). Such structural differences can influence whole-tree water transport/hydraulic conductance and transpiration responses (Simonin et al., 2006). Although the present model scales stand-level maximum xylem conductance with *LAI* (A.1.2.1), it assumes a fixed relationship between leaf area and hydraulic capacity and does not explicitly account for changes in sapwood area or sapwood-specific hydraulic conductivity during canopy recovery.

Similarly, the model has a limited representation of forest root structure, with *RAI* held constant following thinning. In reality, thinning can alter belowground biomass and water uptake. For example, Dore et al. (2010) reported approximately 20% reductions in both coarse-root and fine-root carbon stocks following thinning at US-Fmf. At FI-Hyy, de Quesada (2026) suggested that the temporary loss of deep living roots reduced water uptake from deeper soil layers, contributing to increased soil moisture following thinning.

It should be noted that the elasticity analysis in Section 4.4.1 shows that the model has low sensitivity to both *RAI* and *Z*_*m*_. Therefore, adjusting these parameters to reflect thinning-induced changes may have limited ability to resolve the potential underestimation of hydrological responses to thinning. However, weak sensitivity to these parameters does not imply that root dynamics are ecologically unimportant. Rather, it indicates that the root properties represented by these parameters have limited influence under the current model formulation and site conditions. Capturing additional effects of thinning-induced changes in root structure and function may therefore require modifications to the model formulation.

In this model, leaf area dynamics are represented through a particularly simplified mechanism. The modeled recovery trajectories are strongly influenced by the assumptions underlying the growth formulation. In particular, *g*_+_ is calibrated by assuming that annual foliage production exactly balances litterfall in the unthinned stand, making *LAI*_0_ an equilibrium of the model toward which the simulated trajectories recover without overshooting (Section 3.3.5). The model also assumes a constant annual proportion of foliage loss, although actual litterfall may vary with drought stress, leaf age, and environmental conditions. Together, these simplifications introduce uncertainty into the predicted magnitude and timing of *LAI* recovery, and consequently the duration of thinning-induced changes in soil moisture and water fluxes.

#### 5.5.2 Parameterization and Data Limitations

The present model was tested at only four forest sites spanning mediterranean, temperate, and boreal climates. This limited sample size is primarily due to the scope and time constraints of the study. Thus, although these sites exhibit variation in climatic conditions and stand characteristics (i.e. plant hydraulic traits and *LAI*_0_), it is insufficient to draw broadly generalizable conclusions about ecohydrological responses to thinning across different forest types and climatic conditions. Instead, these simulations serve as an initial exploration of model behavior under contrasting environmental settings. For example, Section 5.2 discusses potential differences in the effects of thinning on *ET* between energy-limited and water-limited forests. With a larger sample of sites representing both conditions, this distinction could be investigated more systematically to determine whether consistent patterns emerge. Furthermore, tropical and deciduous forests are not represented in the present simulations for the reasons discussed in Section 3.3, however, extending the simulations to these forest types could be valuable in evaluating corresponding model behavior.

Moreover, because consistent site-specific field measurements were unavailable, several model parameters were estimated from broader plant-trait compilations or biome-level averages rather than direct measurements at individual sites (Fig.1.3). Thus, more accurate site-specific parameter values could help reduce parameter uncertainty and improve the reliability of model predictions.

## 6. Conclusion

To summarize the results of this study, the major findings and their implications from both ecological and forest management perspectives are presented below:

- Soil saturation responses to forest thinning vary across sites. Although thinning can improve soil water availability, this benefit is not guaranteed, particularly in forests with initially sparse stands where soil evaporation dominates total *ET*.
- Total *ET* remains relatively stable following thinning despite substantial changes in its components, as (1) soil evaporation increases as canopy removal allows more energy to reach the soil surface, offsetting reductions in stand-level transpiration; and (2) improved soil water availability can partially compensate for reduced stand-level transpiration by allowing the remaining foliage to transpire at higher rates.
- Forest thinning can increase leaf-area growth efficiency through alleviated water limitation and reduced self-shading, even as reduced canopy light interception decreases total stand-level leaf-area production.
- Total *ET* is more strongly regulated by precipitation inputs and atmospheric evaporative demand, with their relative importance depending on whether an ecosystem is water or energy-limited. Vegetation characteristics (i.e *LAI* and plant hydraulic traits) have greater effects on *ET* partitioning, as most soil water recharge is eventually lost through either soil evaporation or plant transpiration (or through drainage under sufficiently wet conditions).

Despite its limitations and simplifications, the model demonstrates reasonable performance, with simulated *ET* showing generally good agreement with FLUXNET observations across all four sites. It is also worth noting that although FLUXNET data were used in this study, the model does not necessarily require eddy covariance measurements, as climatic inputs such as precipitation and *E*_0_ can also be obtained from field sensors or remote sensing products. Additionally, stationary climate conditions were assumed in this study to isolate the effects of thinning. However, the modeling framework could also incorporate interannual climate variability by updating both climatic inputs and *LAI* each year with sufficient interannual climate records.

Although the model operates at the stand level, does not explicitly represent spatial heterogeneity in vegetation structure, and has limited capacity for precise quantitative predictions, it provides useful mechanistic insights into canopy–water interactions, soil saturation and *ET* dynamics following thinning, and how these responses vary under different climatic and stand conditions. Its relatively low data requirements and computational simplicity make it a potentially useful low-cost complementary tool alongside field observations and more detailed numerical models, helping address the challenges of long-term field monitoring and informing forest management across a wider range of sites with limited data.

## 7. CRediT author statement

Wandi Feng: Conceptualization, Methodology, Software, Formal analysis, Investigation, Data curation, Visualization, Writing – original draft.

Benjamin Wong Blonder: Conceptualization, Methodology, Supervision, Writing – review & editing.

## Acknowledgments

We thank Dr. Maoya Bassiouni for helping formulate the initial research idea and providing important guidance throughout the project. This research was supported by the University of California, Berkeley Summer Undergraduate Research Fellowship (SURF) program in 2026. Dr. Blonder was supported by the U.S. National Science Foundation under award IOS-2140428.

## Appendix

### A.1 Model Framework

#### A.1.1 Model Inputs

Key inputs to the coupled ecohydrological and leaf-area growth model, including climatic forcing, soil properties, plant hydraulic traits, canopy characteristics, and vegetation growth parameters, as well as model constants, with their corresponding definitions, units, and baseline values (where applicable).

##### A.1.1.1 Climate

| Symbol | Definition | Units/Baseline Value |
|---|---|---|
| *E*<sub>0</sub> | Potential evaporation | *m day*<sup>−1</sup> |
| α | Mean rainfall event depth | *m* |
| λ | Mean rainfall event frequency | *day*<sup>−1</sup> |
| *T*<sub>*g*</sub> | Growing season length | *day* |

##### A.1.1.2 Soil Properties

|  |  |  |
|---|---|---|
| K<sub>s</sub> | Saturated hydraulic conductivity | m day<sup>−1</sup> |
| n | Soil porosity | m<sup>3</sup> m<sup>−3</sup> |
| Z<sub>m</sub> | Rooting zone depth | m |
| Ψ<sub>s0</sub> | Soil water potential near saturation | MPa |
| b | Brooks-Corey pore size exponent | − |
| s<sub>fc</sub> | Soil saturation at field capacity | − |
| s<sub>h</sub> | Soil saturation at hygroscopic point | − |
| f<sub>star</sub> | Transpiration fraction defining incipient stomatal closure threshold s<sup>*</sup> | 0. 95 (−) |
| f<sub>wilt</sub> | Transpiration fraction defining wilting point threshold s<sub>wilt</sub> | 0. 05 (−) |

##### A.1.1.3 Plant Hydraulic Traits

| *k*<sub>*xl,max*</sub> | Leaf-specific maximum xylem conductivity | *kg m*<sup>−1</sup> *MPa*<sup>−1</sup> *s*<sup>−1</sup> |
|---|---|---|
| Ψ<sub>*g*50</sub> | Water potential at 50% stomatal conductance loss | *MPa* |
| Ψ<sub>*x*50</sub> | Water potential at 50% xylem conductance loss | *MPa* |

##### A.1.1.4 Vegetation Structure

|  |  |  |
|---|---|---|
| *LAI*<sub>0</sub> | Initial pre-thinning Leaf Area Index | *m*<sup>2</sup>*m*<sup>−2</sup> |
| *RAI* | Root area index | *m*<sup>2</sup>*m*<sup>−2</sup> |
| *h*<sub>*c*</sub> | Canopy height | *m* |
| *h*<sub>*i*</sub> | Rainfall interception per unit *LAI* per event | 0. 0002 (*m LAI*<sup>−1</sup>*Event*<sup>−1</sup>) |
| *k*<sub>*ext*</sub> | Canopy radiation extinction coefficient | 0. 55 (−) |
| *dr* | Root diameter | 0. 0005 (*m*) |

##### A.1.1.5 Leaf Area Growth

|  |  |  |
|---|---|---|
| *g*<sub>+</sub> | Maximum leaf area growth rate under well-watered conditions | *m*<sup>2</sup>*m*<sup>−2</sup> *day*<sup>−1</sup> |
| ξ | Soil moisture threshold below which growth is reduced | *s*<sup>\*</sup> |
| *ŝ* | Soil moisture threshold below which growth ceases | *s*<sub>*wilt*</sub> |
| *a* | Growth sensitivity exponent to water limitation | 1 (−) |
| β | Annual leaf turnover rate/litterfall fraction | − |

##### A.1.1.6 Other Model Constants

|  |  |  |
|---|---|---|
| ρ<sub>*w*</sub> | Density of liquid water, unit conversion between mass and volume flux | 1000 (*kg m*<sup>−3</sup>) |
| *g* | Gravitational acceleration | 9. 81 (*m s*<sup>−2</sup>) |
| *T*<sub>*d*</sub> | Day length | 43200 (*s day*<sup>−1</sup>) |

#### A.1.2 Model Equations

Model equations used in the coupled ecohydrological and leaf-area growth framework, including hydraulic conductances, soil-water loss functions, soil saturation probability distributions, water flux functions, and leaf-area growth equations, with their corresponding descriptions and units.

##### A.1.2.1 Hydraulic Conductances

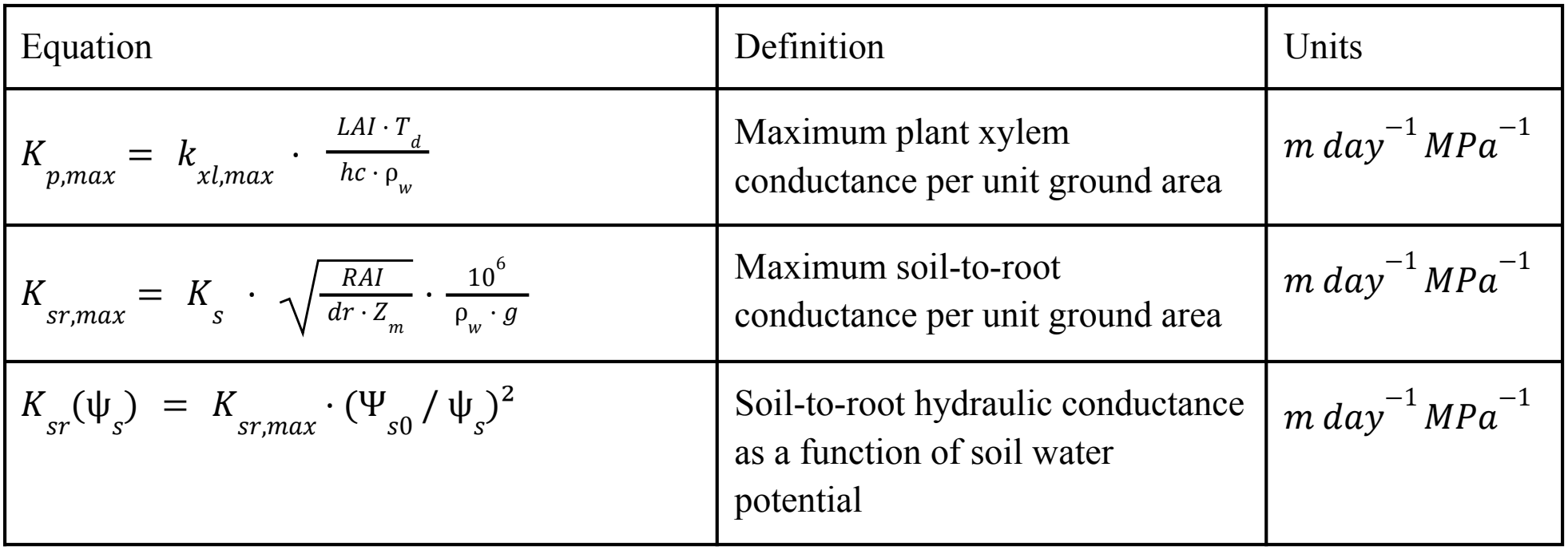

| Equation | Definition | Units |
|---|---|---|
| $K_{p,max} = k_{xl,max} \cdot \frac{LAI \cdot T_d}{hc \cdot \rho_w}$ | Maximum plant xylem conductance per unit ground area | m day<sup>−1</sup> MPa<sup>−1</sup> |
| $K_{sr,max} = K_s \cdot \sqrt{\frac{RAI}{dr \cdot Z_m}} \cdot \frac{10^6}{\rho_w \cdot g}$ | Maximum soil-to-root conductance per unit ground area | m day<sup>−1</sup> MPa<sup>−1</sup> |
| K<sub>sr</sub>(ψ<sub>s</sub>) = K<sub>sr,max</sub> · (Ψ<sub>s0</sub> / ψ<sub>s</sub>)<sup>2</sup> | Soil-to-root hydraulic conductance as a function of soil water potential | m day<sup>−1</sup> MPa<sup>−1</sup> |

##### A.1.2.2 Π Group

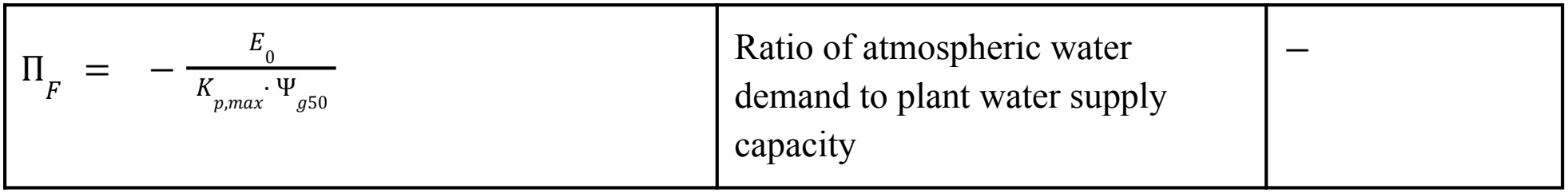
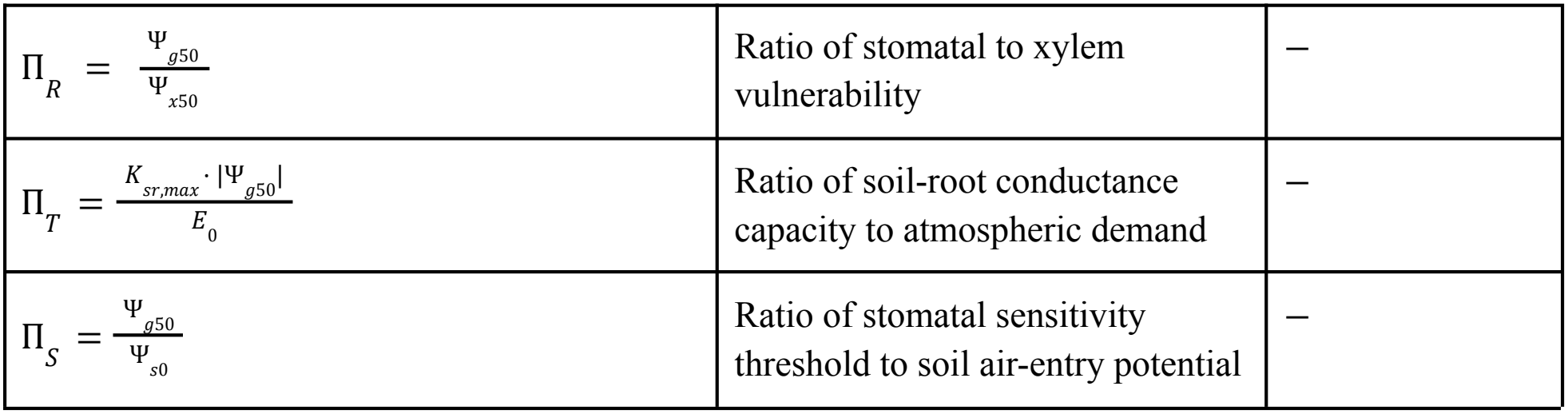

|  |  |  |
|---|---|---|
| $\Pi_{F} = -\frac{E_{0}}{K_{p,max}\cdot\Psi_{g50}}$ | Ratio of atmospheric water demand to plant water supply capacity | − |
| $\Pi_{R} = \frac{\Psi_{g50}}{\Psi_{x50}}$ | Ratio of stomatal to xylem vulnerability | − |
| $\Pi_{T} = \frac{K_{sr,max}\cdot\lvert\Psi_{g50}\rvert}{E_{0}}$ | Ratio of soil-root conductance capacity to atmospheric demand | − |
| $\Pi_{S} = \frac{\Psi_{g50}}{\Psi_{s0}}$ | Ratio of stomatal sensitivity threshold to soil air-entry potential | − |

##### A.1.2.3 Well-Watered Transpiration Ratio and Thresholds

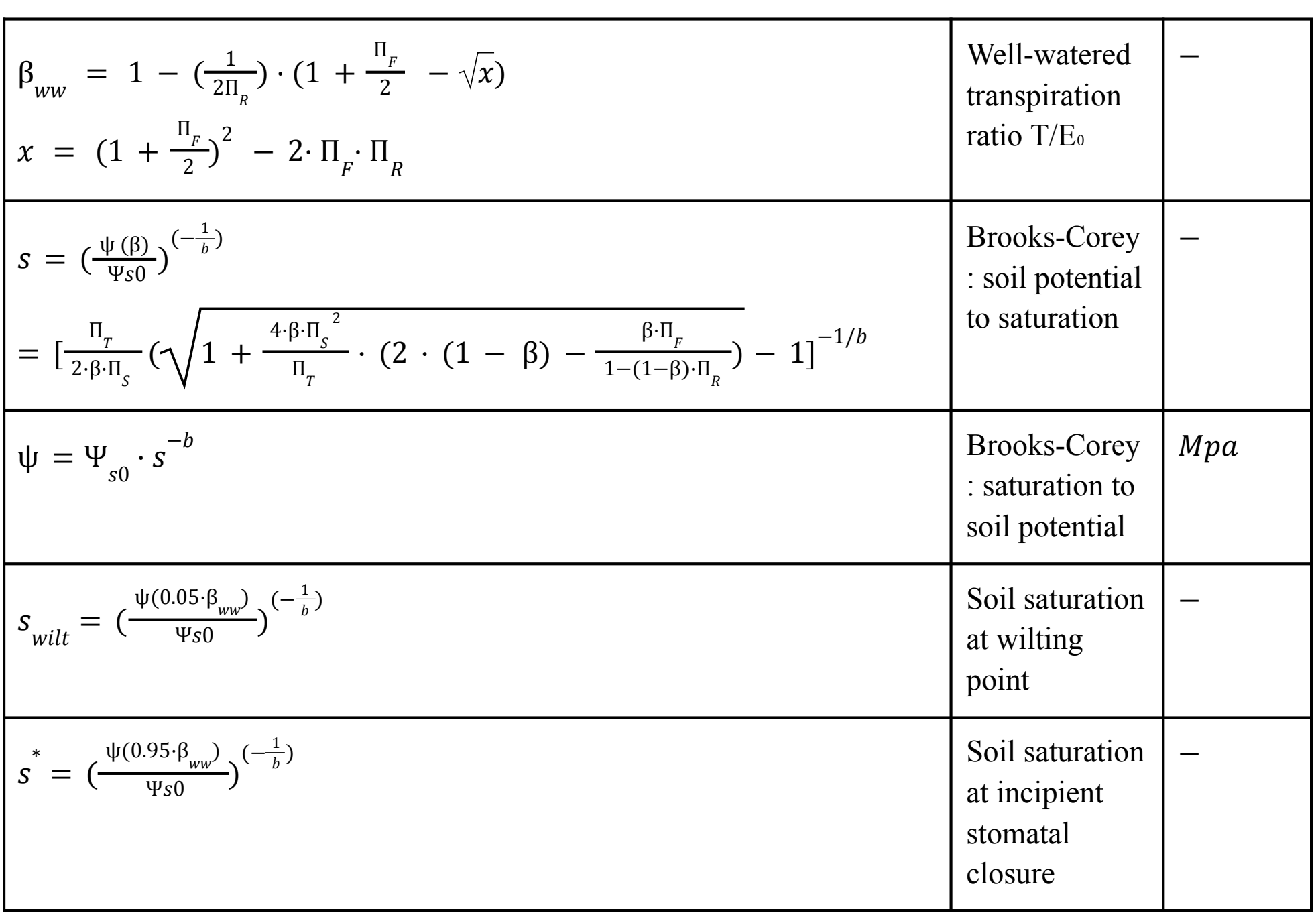

| $\beta_{ww} = 1 - (\frac{1}{2\Pi_{R}}) \cdot (1 + \frac{\Pi_{F}}{2} - \sqrt{x})$<br>$x = (1 + \frac{\Pi_{F}}{2})^{2} - 2 \cdot \Pi_{F} \cdot \Pi_{R}$ | Well-watered transpiration ratio T/E<sub>0</sub> | – |
|---|---|---|
| $s = (\frac{\psi\,(\beta)}{\Psi s0})^{(-\frac{1}{b})}$<br>$= [\frac{\Pi_{T}}{2\cdot\beta\cdot\Pi_{S}}(\sqrt{1 + \frac{4\cdot\beta\cdot\Pi_{S}^{\ 2}}{\Pi_{T}} \cdot (2 \cdot (1 - \beta) - \frac{\beta\cdot\Pi_{F}}{1-(1-\beta)\cdot\Pi_{R}})} - 1]^{-1/b}$ | Brooks-Corey : soil potential to saturation | – |
| $\psi = \Psi_{s0} \cdot s^{-b}$ | Brooks-Corey : saturation to soil potential | *Mpa* |
| $s_{wilt} = (\frac{\psi(0.05\cdot\beta_{ww})}{\Psi s0})^{(-\frac{1}{b})}$ | Soil saturation at wilting point | – |
| $s^{*} = (\frac{\psi(0.95\cdot\beta_{ww})}{\Psi s0})^{(-\frac{1}{b})}$ | Soil saturation at incipient stomatal closure | – |

##### A.1.2.4 Rainfall and Interception

|  |  |  |
|---|---|---|
| $\delta = h_{i} \cdot LAI$ | Total canopy interception per rainfall event | *m* |
| $\lambda^{*} = \lambda \cdot exp(-\frac{\delta}{\alpha})$ | Effective rainfall frequency reaching soil | *day*<sup>−1</sup> |
| $\phi = e^{-k_{ext} \cdot LAI}$ | Fraction of radiation reaching soil surface | − |

##### A.1.2.5 Soil Water Loss Function

| $E_{max} = \frac{\phi \cdot E_0}{n \cdot Z_m}$ | Normalised maximum soil evaporation rate | $day^{-1}$ |
|---|---|---|
| $T_{ww} = \frac{(1 - \phi) \cdot \beta_{ww} \cdot E_0}{(n \cdot Z_m)}$ | Normalised well-watered transpiration rate | $day^{-1}$ |
| $bb = 2b + 4$ | Brooks-Corey drainage exponent | − |
| $m = \frac{K_s}{n \cdot Z_m \cdot ( e^{bb \cdot (1 - s_{fc})} - 1)}$ | Normalized drainage coefficient | $day^{-1}$ |
| $s > s_{fc}$:<br>$\rho(s) = E_{max} + T_{ww} + m \cdot (e^{bb \cdot (s - s_{fc})} - 1)$<br>$s^{*} < s \leq s_{fc}$:<br>$\rho(s) = \frac{E_{max} \cdot (s - s_h)}{(s_{fc} - s_h)} + T_{ww}$<br>$s_w < s \leq s^{*}$:<br>$\rho(s) = \frac{E_{max} \cdot (s - s_h)}{(s_{fc} - s_h)} + \frac{T_{ww} \cdot (s - s_{wilt})}{(s^{*} - s_{wilt})}$<br>$s_h < s \leq s_{wilt}$:<br>$\rho(s) = \frac{E_{max} \cdot (s - s_h)}{(s_{fc} - s_h)}$<br>$s \leq s_h$:<br>$\rho(s) = 0$ | Loss function | $day^{-1}$ |

##### A.1.2.6 Soil Saturation PDF

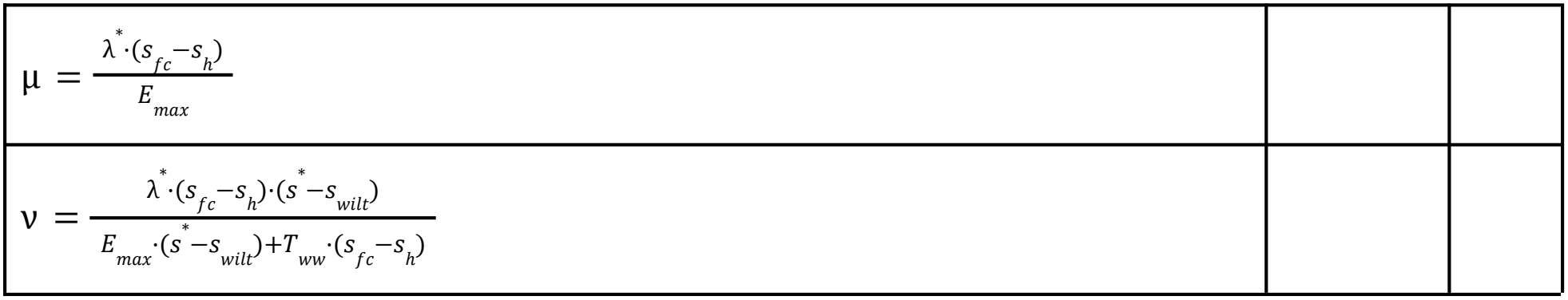
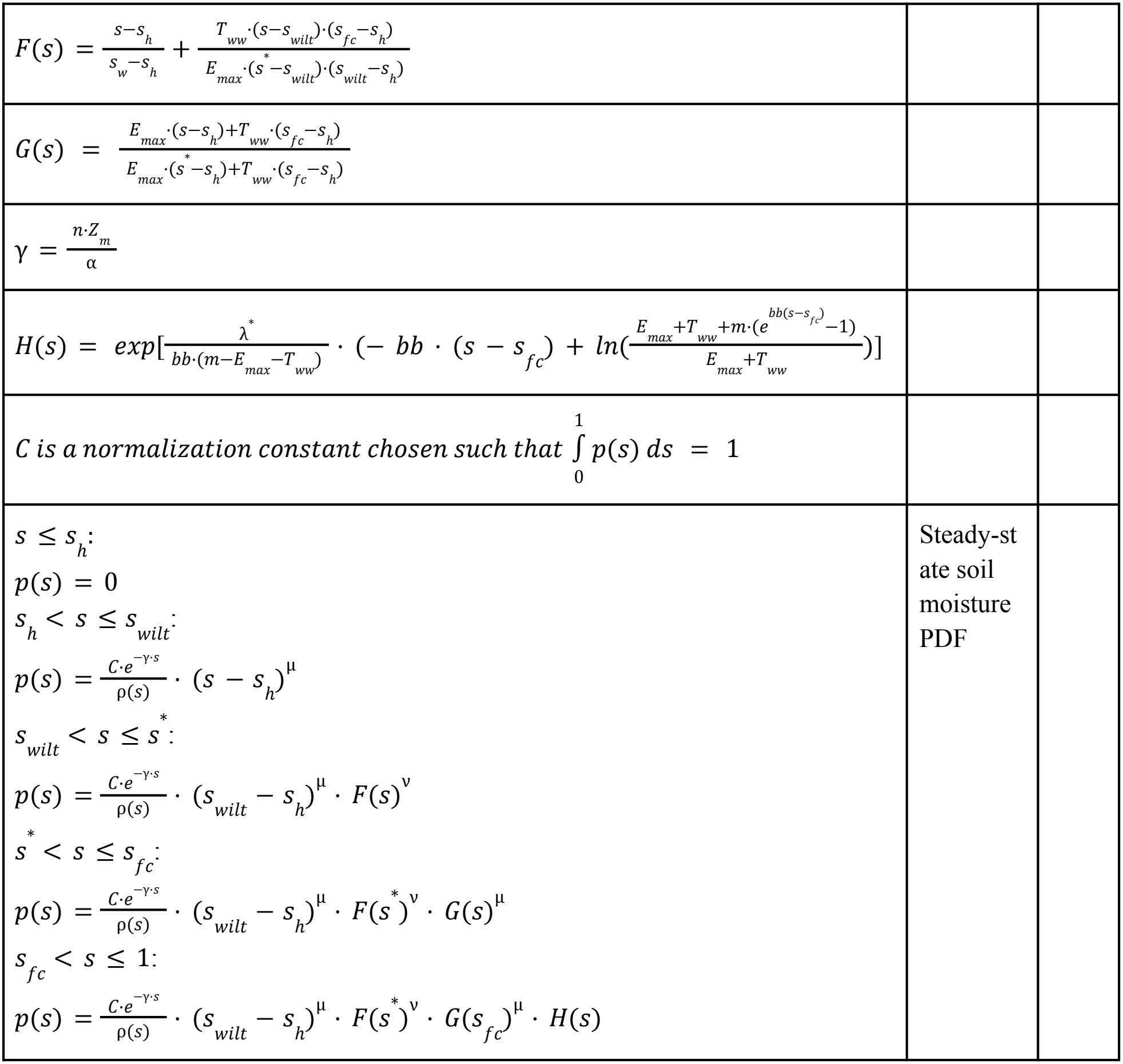

|  |  |  |
|---|---|---|
| $\mu = \frac{\lambda^{*}\cdot(s_{fc}-s_{h})}{E_{max}}$ |  |  |
| $\nu = \frac{\lambda^{*}\cdot(s_{fc}-s_{h})\cdot(s^{*}-s_{wilt})}{E_{max}\cdot(s^{*}-s_{wilt})+T_{ww}\cdot(s_{fc}-s_{h})}$ |  |  |
| $F(s) = \frac{s-s_{h}}{s_{w}-s_{h}} + \frac{T_{ww}\cdot(s-s_{wilt})\cdot(s_{fc}-s_{h})}{E_{max}\cdot(s^{*}-s_{wilt})\cdot(s_{wilt}-s_{h})}$ |  |  |
| $G(s) = \frac{E_{max}\cdot(s-s_{h})+T_{ww}\cdot(s_{fc}-s_{h})}{E_{max}\cdot(s^{*}-s_{h})+T_{ww}\cdot(s_{fc}-s_{h})}$ |  |  |
| $\gamma = \frac{n\cdot Z_{m}}{\alpha}$ |  |  |
| $H(s) = exp[\frac{\lambda^{*}}{bb\cdot(m-E_{max}-T_{ww})}\cdot(- bb\cdot(s-s_{fc}) + ln(\frac{E_{max}+T_{ww}+m\cdot(e^{bb(s-s_{fc})}-1)}{E_{max}+T_{ww}}))]$ |  |  |
| $C\ is\ a\ normalization\ constant\ chosen\ such\ that\ \int_{0}^{1} p(s)\,ds = 1$ |  |  |
| $s \leq s_{h}$:<br>$p(s) = 0$<br>$s_{h} < s \leq s_{wilt}$:<br>$p(s) = \frac{C\cdot e^{-\gamma\cdot s}}{\rho(s)}\cdot(s-s_{h})^{\mu}$<br>$s_{wilt} < s \leq s^{*}$:<br>$p(s) = \frac{C\cdot e^{-\gamma\cdot s}}{\rho(s)}\cdot(s_{wilt}-s_{h})^{\mu}\cdot F(s)^{\nu}$<br>$s^{*} < s \leq s_{fc}$:<br>$p(s) = \frac{C\cdot e^{-\gamma\cdot s}}{\rho(s)}\cdot(s_{wilt}-s_{h})^{\mu}\cdot F(s^{*})^{\nu}\cdot G(s)^{\mu}$<br>$s_{fc} < s \leq 1$:<br>$p(s) = \frac{C\cdot e^{-\gamma\cdot s}}{\rho(s)}\cdot(s_{wilt}-s_{h})^{\mu}\cdot F(s^{*})^{\nu}\cdot G(s_{fc})^{\mu}\cdot H(s)$ | Steady-state soil moisture PDF |  |

##### A.1.2.7 Mean Water Fluxes

|  |  |  |
|---|---|---|
| $< E > = \int_{0}^{1} E_{loss}(s) \cdot p(s) \cdot ds$<br>$= \phi \cdot E_0 \int_{s_h}^{s_{fc}} \frac{s-s_h}{s_{fc}-s_h} \cdot p(s) \cdot ds + \phi \cdot E_0 \int_{s_{fc}}^{1} p(s) \cdot ds$ | Mean growing-season evaporation | $m\, day^{-1}$ |
| $< T > = \int_{0}^{1} T_{loss}(s) \cdot p(s) \cdot ds$ | Mean growing-season transpiration | $m\, day^{-1}$ |
| $= (1 - \phi) \cdot \beta_{ww} \cdot E_0 \int_{s_w}^{s^*} \frac{s-s_w}{s^*-s_w} p(s)ds + (1 - \phi) \cdot \beta_{ww} \cdot E_0 \int_{s^*}^{1} p(s)ds$ |  |  |
| $< ET > = < T > + < E >$ | Mean growing-season evapotranspiration | $m\, day^{-1}$ |
| $< I > = \alpha \cdot \lambda \cdot (1 - e^{-\frac{\delta}{\alpha}})$ | Mean growing-season interception | $m\, day^{-1}$ |

##### A.1.2.8 Leaf Area Growth

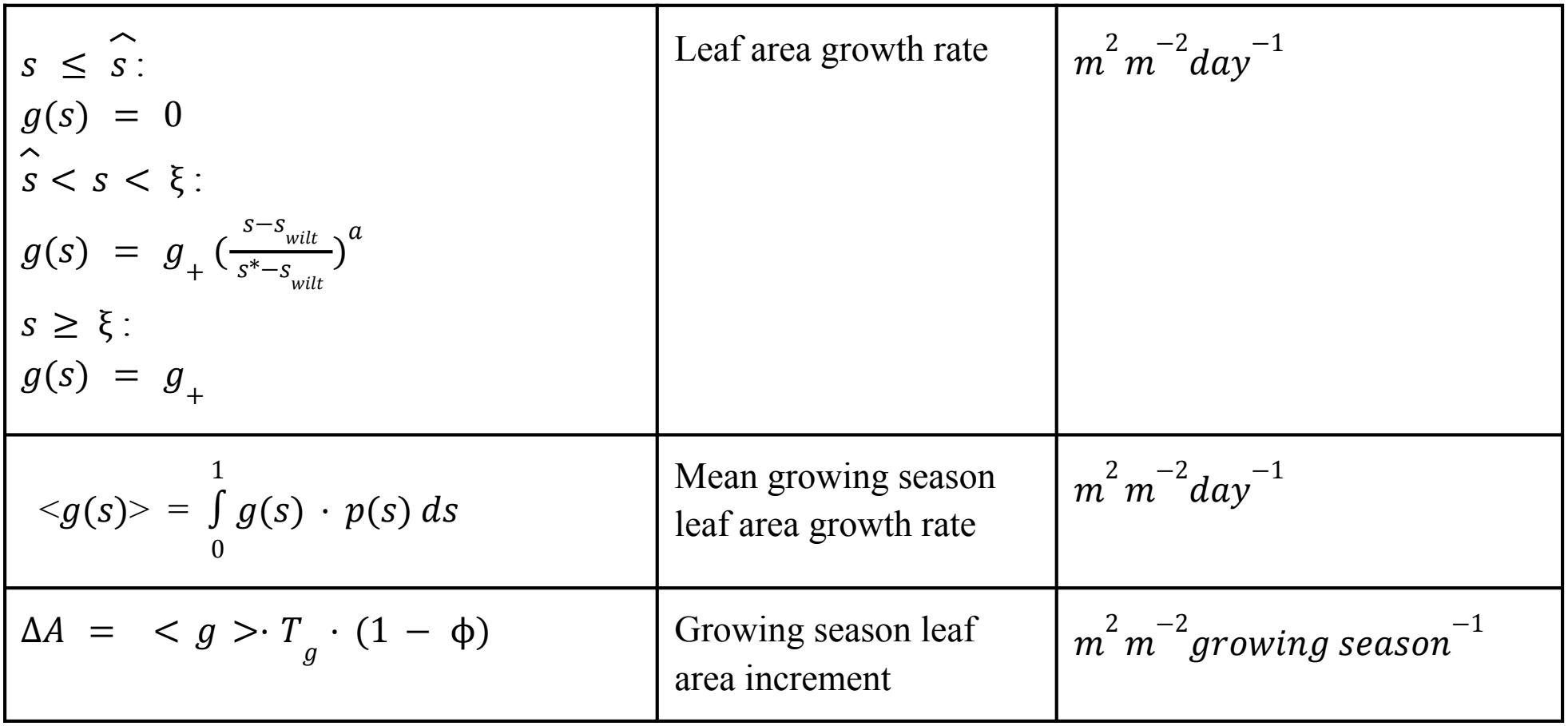

|  |  |  |
|---|---|---|
| $s \leq \hat{s}$:<br>$g(s) = 0$<br>$\hat{s} < s < \xi$:<br>$g(s) = g_{+}(\frac{s-s_{wilt}}{s^{*}-s_{wilt}})^{a}$<br>$s \geq \xi$:<br>$g(s) = g_{+}$ | Leaf area growth rate | m<sup>2</sup>m<sup>−2</sup>day<sup>−1</sup> |
| $<g(s)> = \int_{0}^{1} g(s) \cdot p(s)\, ds$ | Mean growing season leaf area growth rate | m<sup>2</sup>m<sup>−2</sup>day<sup>−1</sup> |
| ΔA = < g >· T<sub>g</sub> · (1 − ϕ) | Growing season leaf area increment | m<sup>2</sup>m<sup>−2</sup>growing season<sup>−1</sup> |

### A.2 Simulation Inputs

Model inputs for the four simulated forest sites (US-Fmf, CA-TP4, FR-Pue, and FI-Hyy), including climatic parameters, soil properties, plant hydraulic traits, canopy characteristics, and vegetation growth parameters, with their corresponding values, units, and literature sources where applicable.

|  | US-Fmf | CA-TP4 | FR-Pue | FI-Hyy |
|---|---|---|---|---|
| *T*<sub>*g*</sub><br>(*days*) | 109 (May 15 - Aug 31)<br>(Teets et al., 2026) | 109 (May 15 - Aug 31)<br>(Wang & Chen, 2012) | 106 (Apr 01 - July 15)<br>( Limousin et al., 2012; Ruffault et al., 2023) | 78 (May 15 - July 31)<br>(Rautiainen and Lukeš, 2015; Launiainen, 2010) |
| Growing Season λ<br>(*day*<sup>−1</sup>) | 0.3440 | 0.5304 | 0.2683 | 0.5009 |
| Growing Season α<br>(*m event*<sup>−1</sup>) | 0.0065 | 0.0063 | 0.0073 | 0.0051 |
| Growing Season average *E*<sub>0</sub><br>(*m day*<sup>−1</sup>) | 0.0048 | 0.0046 | 0.0043 | 0.0035 |
| *h*<sub>*c*</sub><br>(*m*) | 18<br>(Dore & Kolb, 2026) | 21<br>(Arain, 2016) | 5<br>(Ourcival et al., 2016) | 20<br>(Mammarella et al., 2020) |
| ψ<sub>*g*50</sub><br>(*MPa*) | -1.44 | -1.44 | -1.46 | -1.89 |
| ψ<sub>*x*50</sub><br>(*MPa*) | -4.88 | -4.88 | -5.46 | -3.68 |
| *k*<sub>*xl,max*</sub><br>(*kg m*<sup>−1</sup> *MPa*<sup>−1</sup> *s*<sup>−1</sup>) | 0.00032 | 0.00032 | 0.00035 | 0.00040 |
| Soil Texture | 3 | 6 | 1 | 4 |
|  | (Dore et al., 2010) | (Khomik et al., 2010) | (Cabon et al., 2018) | (Churkina et al., 2003) |
| *RAI*<br>(*m*<sup>2</sup>*m*<sup>−2</sup>) | 11 | 11 | 11.6 | 4.6 |
| β | 0.25<br>(Insinna et al., 2007) | 0.4<br>(Wang and Chen, 2012) | 0.35<br>( Limousin et al., 2012; Terradas, 1999) | 0.25<br>(Launiainen, 2010) |
| *LAI*<sub>0</sub><br>(*m*<sup>2</sup>*m*<sup>−2</sup>) | 1.5<br>(Dore et al., 2010) | 5<br>(Wang and Chen, 2012; Khomik et al., 2010) | 2.2<br>(Cabon et al., 2018; Limousin et al., 2012; Ruffault et al., 2023) | 2.2<br>(Peräkylä et al., 2025; Rautiainen and Lukeš, 2015) |

#### A.2.1 Soil Texture Classification

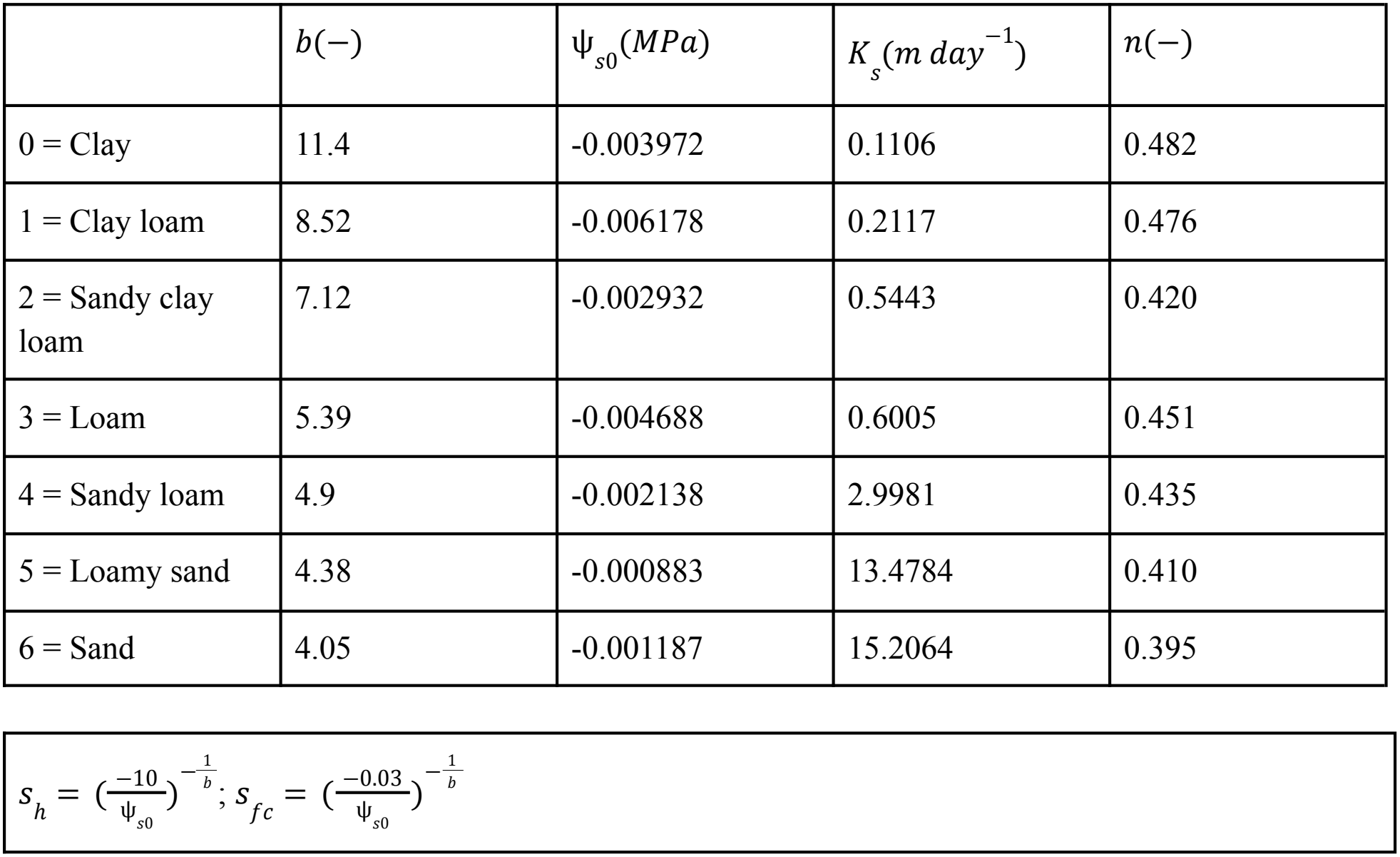

|  | b(−) | ψ<sub>s0</sub>(MPa) | K<sub>s</sub>(m day<sup>−1</sup>) | n(−) |
|---|---|---|---|---|
| 0 = Clay | 11.4 | -0.003972 | 0.1106 | 0.482 |
| 1 = Clay loam | 8.52 | -0.006178 | 0.2117 | 0.476 |
| 2 = Sandy clay loam | 7.12 | -0.002932 | 0.5443 | 0.420 |
| 3 = Loam | 5.39 | -0.004688 | 0.6005 | 0.451 |
| 4 = Sandy loam | 4.9 | -0.002138 | 2.9981 | 0.435 |
| 5 = Loamy sand | 4.38 | -0.000883 | 13.4784 | 0.410 |
| 6 = Sand | 4.05 | -0.001187 | 15.2064 | 0.395 |

$s_{h} = (\frac{-10}{\psi_{s0}})^{-\frac{1}{b}}; s_{fc} = (\frac{-0.03}{\psi_{s0}})^{-\frac{1}{b}}$

#### A.3 Codes and data availability

All the scripts/data required to reproduce the results of this study are available at DOI: 10.5281/zenodo.24071679. To replicate the study, follow the instructions in README.md.

Illustration graphs were created in https://BioRender.com

